# Canonical Wnt pathway modulation is required to correctly execute multiple independent cellular dynamic programs during cranial neural tube closure

**DOI:** 10.1101/2024.12.19.629501

**Authors:** Amber Huffine Bogart, Eric R. Brooks

**Affiliations:** Department of Molecular Biomedical Sciences, College of Veterinary Medicine, North Carolina State University

**Keywords:** Wnt, Lrp6, Apc, neural tube closure, apical constriction, proliferation

## Abstract

Defects in cranial neural tube closure are among the most common and deleterious human structural birth defects. Correct cranial closure requires the coordination of multiple cell dynamic programs including cell proliferation and cell shape change. Mutations that impact Wnt signaling, including loss of the pathway co-receptor LRP6, lead to defects in cranial neural tube closure, but the cellular dynamics under control of the Wnt pathway during this critical morphogenetic process remain unclear. Here, we use mice mutant for LRP6 to examine the consequences of conditional and global reduction in Wnt signaling and mutants with conditional inactivation of APC to examine the consequences of pathway hyperactivation. Strikingly, we find that regulated Wnt signaling is required for two independent events during cranial neural tube closure. First, global reduction of Wnt leads to a surprising hyperplasia of the cranial neural folds driven by excessive cell proliferation at early pre-elevation stages, with the increased tissue volume creating a mechanical blockade to efficient closure despite normal apical constriction and cell polarization at later stages. Conversely, conditional hyperactivation of the pathway at later elevation stages prevents correct actin organization, blocking apical constriction and neural fold elevation without impacting tissue scaling. Together these data reveal that Wnt signaling levels must be modulated to restrict proliferation at early stages and promote apical constriction at later elevation stages to drive efficient closure of the cranial neural tube.

## Introduction

Formation of the brain requires the conversion of the cranial neural plate, a sheet of neuroepithelial cells, into the closed tube that will form the structural basis of the nervous system. This process is known as cranial neural tube closure and defects in its execution are among the most common and deleterious human birth defects, resulting in invariable pre- or perinatal lethality (Copp et al., 2013; Wallingford et al., 2013; Wilde et al., 2014; Zaganjor et al., 2016). Cranial closure exhibits complex genetic requirements, with well over a hundred genes implicated in the process (Harris and Juriloff, 2007; Harris and Juriloff, 2010; Lee and Gleeson, 2020; Wilde et al., 2014); however, the specific function of the majority of these genes in cranial tissue remodeling remains unknown.

The closure process (schematized in Figure 1A) begins with the induction of the anterior neural tissues, which completes by day 7.5 (E7.5) in the mouse embryo (Harland, 2000; Levine and Brivanlou, 2007; Ozair et al., 2013; Rhinn et al., 1998). The cranial neural plate continuously grows via proliferation during and after induction, resulting in the formation of a wide, outwardly curved morphology, with the lateral edges of the tissue situated below the level of more medial regions (Jacobson and Tam, 1982; Juriloff and Harris, 2018; Morriss-Kay, 1981). Beginning at E8, the tissue undergoes a curvature inversion that will bring these lateral edges above the midline, a process known as neural fold elevation. Subsequently, the edges will deflect inward to meet above the midline of the tissue during apposition. Finally, the neural folds will fuse, as will the overlying surface ectoderm, completing closure and internalizing the tissue.

**Figure 1:**
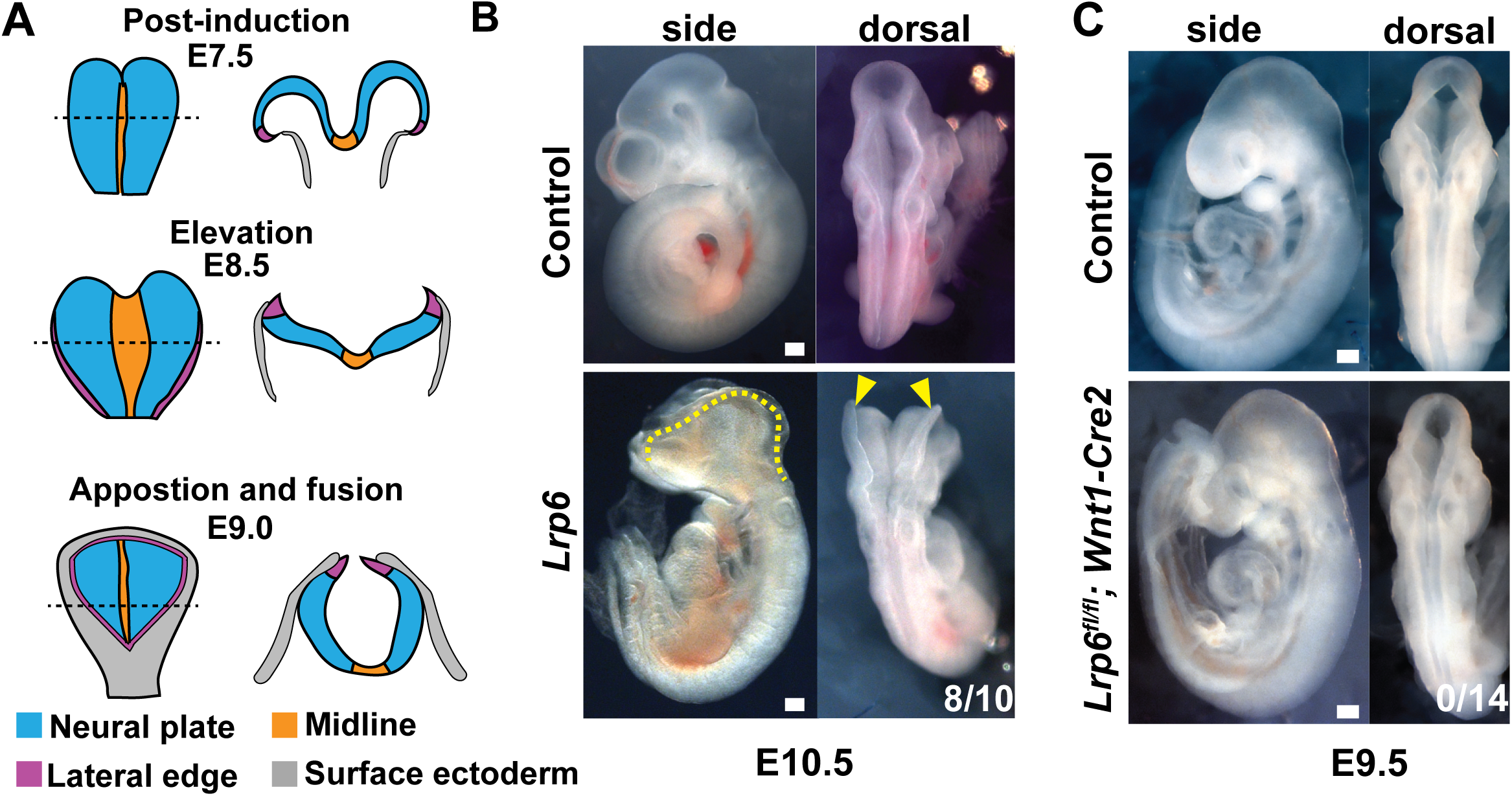
Cranial closure defects occur in global but not cranial neural fold specific *Lrp6* mutants. (**A**) Schematics representing the *en face* (left) and transverse (right) morphology of the midbrain region of the cranial neural tissues at post-induction (top), elevation (middle) and apposition (bottom) stages of closure. (**B**) Side and dorsal views of control and *Lrp6* global mutants. Mutants exhibit failed cranial closure from the forebrain to the hindbrain-spinal cord juncture (dashed yellow line) at 80% penetrance. Notably, however, the neural folds appear well elevated in these mutants (yellow arrowheads). (**C**) Side and dorsal views of a control embryo and an embryo where *Lrp6* function has been conditionally ablated specifically in the future midbrain tissues at the onset of neural fold elevation via the activity of the *Wnt1-Cre2* transgene. No recovered embryos of this genotype displayed cranial neural tube closure defects. Scale bar, 100 µm; yellow arrowheads show the position of the lateral edge of the cranial neural tissues.

The tissue scale morphogenetic events of cranial closure require dynamic activities of individual cells, including shape changes such as apical constriction to drive the curvature change of neural fold elevation, cell rearrangement to alter the aspect ratio of the tissue, and polarized cell division which is thought to drive anisotropic growth along the anteroposterior axis (Devenport, 2016; Juriloff and Harris, 2018; Nikolopoulou et al., 2017; Vijayraghavan and Davidson, 2017; Wilde et al., 2014). These individual cellular activities must be coordinated in space and time to effectively reshape the tissue but the upstream controls on their spatiotemporal deployment often remain opaque. Developmental morphogens such as Wnt and Shh are likely to be key players in ensuring that cell dynamics are correctly integrated over time and tissue length given that alterations to the activity and/or reach of these pathways lead to gross defects in neural tube closure (Allache et al., 2014; Brooks et al., 2020; Brown et al., 2020; Carter et al., 2005; Gray et al., 2013; Harris and Juriloff, 2007; Harris and Juriloff, 2010; Kimura-Yoshida et al., 2015; Misra and Matise, 2010; Patterson et al., 2009; Pinson et al., 2000; Ybot-Gonzalez et al., 2002). Curiously, both mutations that inhibit or hyperactivate the activity of the Wnt pathway lead to defects in cranial closure (Brown et al., 2020; Gray et al., 2013; Kimura-Yoshida et al., 2015; Pinson et al., 2000). However, the impact of Wnt dysregulation on the spatial and temporal patterns of cell behavior required for closure are not fully understood. This confusion arises, at least in part, from the relative complexity of Wnt signaling in the tissue, where multiple Wnt ligands and receptors are expressed (Bally-Cuif et al., 1995; Bouillet et al., 1996; Parr et al., 1993; Yamaguchi et al., 1999), as well as from the fact that the pathway can exert control over multiple classes of cellular dynamics including proliferation, cell shape change, and planar polarized rearrangement (Butler and Wallingford, 2017; Fumoto et al., 2017; Reya and Clevers, 2005; Steinhart and Angers, 2018; Sutherland et al., 2020; Yoon et al., 2023). Thus, the specific roles of Wnt signaling in the tissue, and whether loss and hyperactivation of the pathway lead to equivalent downstream defects, remain unknown.

Here, we seek to systematically address the function of Wnt signaling during cranial neural tube closure. To do so we use a conditional mutant of the critical pathway co-receptor *Lrp6* to reduce Wnt signaling either globally, or specifically within the future midbrain tissues at the beginning of neural fold elevation. This analysis reveals that Wnt inhibition, surprisingly, does not impact the events of neural fold elevation and that apical constriction and planar polarization are normal. Closure defects in these mutants instead stem from neural fold hyperplasia caused by excessive proliferation just after the completion of anterior neural induction, revealing an entirely novel cellular basis for cranial closure defects. We next ask if hyperactivation of Wnt signaling also leads to defects in closure by conditionally inactivating the critical pathway regulatory factor APC. Unlike *Lrp6* mutants, loss of *Apc* does not alter tissue scale, but instead leads to a dysregulation of actin organization, preventing apical constriction and blocking neural fold elevation. Thus, both reduction and hyperactivation of Wnt activity result in failures in cranial neural tube closure but do so by interfering with distinct cellular mechanisms. Together, our data demonstrate that Wnt signaling levels are key determinants of the success of multiple independent patterns of cell behavior required for robust cranial neural tube closure.

## Results

### Global, but not elevation stage specific, reduction of Wnt signaling leads to failure of cranial neural tube closure

*Lrp6* encodes a critical Wnt pathway co-receptor broadly required for signal transduction, and loss of function mutations in this gene lead to cranial closure defects (Andersson et al., 2010; Gray et al., 2013; Pinson et al., 2000; Ren et al., 2021; Zhao et al., 2022; Zhou et al., 2010). Because *Lrp6* is widely expressed throughout the epiblast, and uniformly in the later cranial neural plate (Geng et al., 2023), we used a conditional allele of *Lrp6* (*Lrp6^fl^* (Joeng et al., 2011)) to explore the temporal requirements for Wnt signaling during closure. First, we used *Sox2-Cre* to ablate *Lrp6* function throughout the epiblast (Hayashi et al., 2002) and to drive germline recombination in order to create a heritable global loss of function allele (*Lrp6^Δ^*). In all analyses reported below no difference was observed between embryos with epiblast specific ablation (*Lrp6 ^Δ/fl^*; *Sox2-Cre)* and those with global germline ablation (*Lrp6 ^Δ/Δ^*), indicating that phenotypes arise from the epiblast. We therefore refer to these genotypes collectively as *Lrp6* mutants. Next, we directly tested the requirement for Wnt activity during neural fold elevation by ablating *Lrp6* function specifically within the future midbrain at the onset of elevation (∼E8.25) using *Wnt1-Cre2* (Chen et al., 2017; Lewis et al., 2013).

Global *Lrp6* loss led to exencephaly with an 80% penetrance (8/10 mutants, Figure 1B), in line with though modestly higher than observed in previously reported *Lrp6* mutants (Gray et al., 2010; Gray et al., 2013; Pinson et al., 2000; Zhou et al., 2010). Notably, despite the clear exencephalic phenotypes, the cranial neural folds appeared well elevated in these global loss of function mutants. Conversely, none of the conditional *Lrp6^fl/fl^; Wnt1-Cre2* mutant embryos we recovered showed failures in cranial closure (Figure 1C, 0/14 mutants; penetrance difference from global *Lrp6* mutants p < 0.0001 by Fisher’s Exact Test, Table S1). Together, these data demonstrate that LRP6 function is broadly required for cranial neural tube closure, and loss of function specifically in the midbrain during neural fold elevation stages is not sufficient to drive exencephaly. We therefore focused the rest of our analyses on global *Lrp6* mutants.

We next wanted to map the consequences of loss of *Lrp6* on the spatial pattern of Wnt signal transduction in the cranial neural plate. The reduction of Wnt signaling in these mutants is expected to result in increased activity of the cytoplasmic destruction complex for β-catenin, preventing its accumulation and the resulting activation of pathway targets (Steinhart and Angers, 2018). However, we were not able to detect any changes in the relative ratio of β-catenin between the junctional and cytoplasmic domains either apically or basally at these stages (Supplemental Figure 1A-F), suggesting either that β-catenin organization does not change in *Lrp6* mutants or, more likely, that such changes are below the detection threshold of this approach. We therefore turned to a previously described transgenic reporter of Wnt activity comprised of H2B-GFP under the control of Tcf/Lef binding sites (Ferrer-Vaquer et al., 2010). In cranial neural plates of control embryos at elevation stages (E8.5), this reporter recapitulated the expected pattern of Wnt signaling (Brown et al., 2020), showing high expression at the lateral edges of the tissue, and a midbrain specific stripe of expression in the WNT1 domain (Figure 2A). *Lrp6* mutant embryos retained a similar spatial pattern of Wnt response but demonstrated apparent diminishment of reporter activity at all positions. To better understand this diminishment, we quantified the relative Wnt signaling strength at progressive distances from the tissue midline specifically within the midbrain stripe. In wild-type control embryos this analysis revealed a graded Wnt response, with the highest levels of signaling observed at the lateral tissue border (Figure 2B, C); in *Lrp6* mutants, pathway response was significantly decreased at several mediolateral positions, mostly within the lateral domain where cells undergo apical constriction (Brooks et al., 2020). Together, these data indicate that *Lrp6* mutants show a diminishment, but not elimination, of Wnt signaling during cranial neural fold elevation.

**Figure 2:**
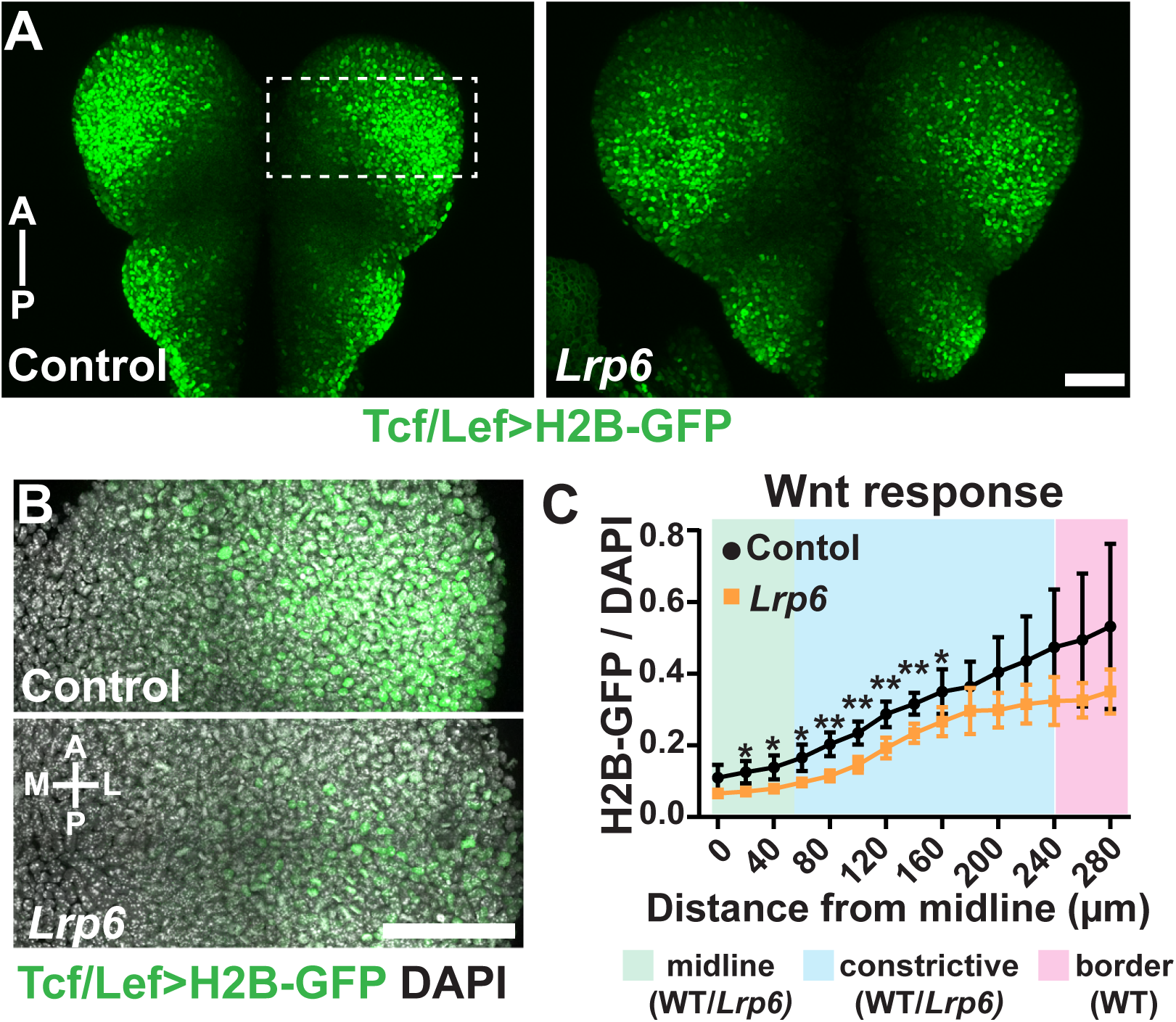
L*r*p6 mutants show reduced Wnt signaling in the midbrain during neural fold elevation. (**A**) Maximum intensity projections of control and *Lrp6* embryos expressing the Tcf/Lef>H2B-GFP reporter of Wnt signaling. The control embryo demonstrates the spatial pattern of Wnt response in the midbrain and hindbrain tissues of the cranial neural plate. Pathway response is high laterally at all anteroposterior positions, with expansion medially occurring specifically in the future midbrain. The gross pattern of Wnt response is not altered in *Lrp6* mutants but shows apparent diminishment in all domains. Dashed box represents the region analyzed in B-C. (**B**) The midbrain stripe of Wnt response in the right neural fold, with DAPI as a counterstain, is shown in control (top) and *Lrp6* mutant (bottom) embryos. (**C**) Quantification of the Wnt signaling response, after normalization to DAPI, is shown at increasing distances from the midline. The positions of the midline, lateral constrictive, and WT border domains are color-coded. Both control and *Lrp6* mutant embryos show increasing response in more lateral domains, however mutants show a significant reduction in response at several positions along the mediolateral axis (two-way ANOVA with Tukey’s multiple comparisons, n = 5 control embryos, 3 *Lrp6* embryos, see Table S1 for p values). Anterior is up in all images, scale bars represent 100 µm. *, p < 0.05, **, p < 0.01.

### *Lrp6* mutants show tissue scaling defects despite normal cell remodeling

To better understand what tissue-level mechanisms could be driving terminal closure defects, we examined *Lrp6* mutants at stages ranging from E8.0-E8.5 when the neural folds are undergoing critical morphogenetic events including anteroposterior growth and elongation, as well as the curvature changes associated with neural fold elevation (Figure 1A). At mid-elevation stages (6- 7 somites, E8.5), *Lrp6* mutants showed a striking tissue scaling defect of the cranial neural plate (Figure 3A-D). The tissue width—defined as the largest mediolateral extent in midbrain tissues— was doubled compared to control littermates. Conversely, tissue length—defined as the distance from the anterior neural ridge to the otic sulcus—was comparable to controls (Figure 3D). This same tissue scaling phenotype was apparent, though less pronounced, at pre-elevation stages (E8.0, 0-3 somites, Figure 3B). To our knowledge this is the first report of cranial neural fold scaling defects at these stages, potentially representing a unique tissue-level etiology for closure defects.

**Figure 3:**
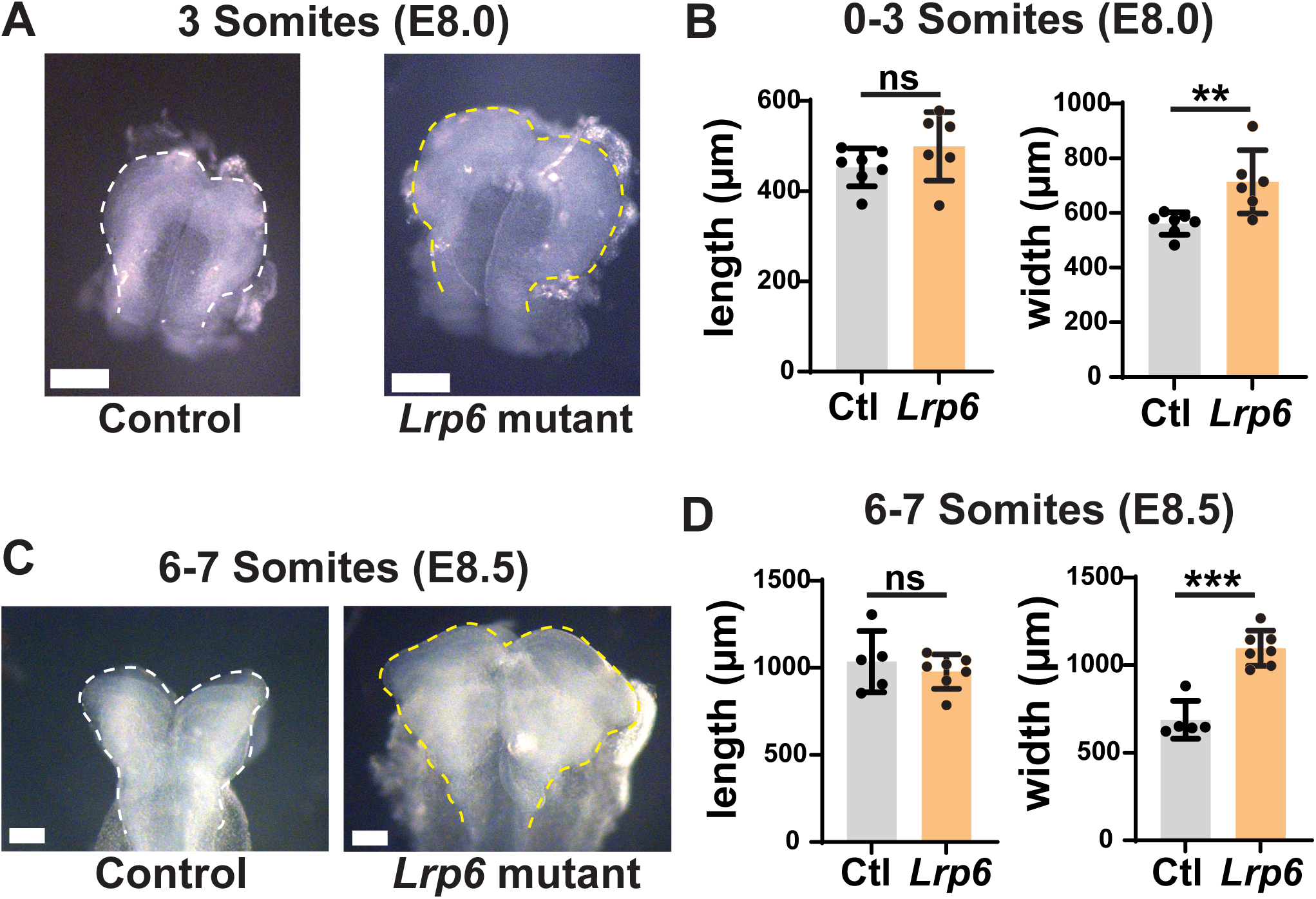
*Lrp6* mutants show enlargement of the cranial neural plate. (**A**) Brightfield images of cranial neural plates from control and *Lrp6* mutant at E8 (3 somites), a stage prior to the onset of strong cranial neural fold elevation. Note that a visible increase in tissue size is already apparent at these stages (compare white dashed line in the control to yellow dashed line in the mutant). The apparent lengthening of *Lrp6* tissue is an artifact caused by delay in formation of the cranial flexure between the forebrain and midbrain tissues. (**B**) Quantification of the length and width of the cranial neural plate in control and *Lrp6* embryos at 0-3 somite stages. Length is defined as the distance between the anterior neural ridge and the otic sulcus. There is no significant difference in this value between control and *Lrp6* mutants (control, mean ± SD: 452.7 ± 41.87 µm; *Lrp6*: 499.1 ± 76.00 µm. n = 7 control and 6 *Lrp6* embryos, p = 0.1905 by unpaired t-test). Width is defined as the length of a line drawn from one lateral edge of the tissue to the other across the widest portion of the future midbrain. This value is significantly larger in *Lrp6* mutans (control, 561.4 ± 40.86 µm; *Lrp6*: 713.9.1 ± 115.6.00 µm. n = 7 control and 6 *Lrp6* embryos, p = 0.0073 by unpaired t-test). (**C-D**) Tissue enlargement in *Lrp6* mutants persists into elevation stages (E8.5, 6-7 somites) and continues to be driven by tissue widening (control, 688.6 ± 108.3 µm; *Lrp6*: 1097 ± 100.9 µm. n = 5 control and 7 *Lrp6* embryos, p < 0.0001 by unpaired t-test) but not lengthening (control, 1035 ± 175.7 µm; *Lrp6*: 978.0 ± 98.83 µm. n = 5 control and 7 *Lrp6* embryos, p = 0.4845 by unpaired t-test). Scale bars represent 100 µm, dashed outlines indicate edges of the neural plate. ns, not significant; **, p < 0.01, ***, p < 0.001. Anterior is up in all images.

We next considered potential cellular mechanisms that could increase the width of the tissue. First, apical constriction reduces the apical surface area of the tissue to drive the curvature changes required for neural fold elevation (Brooks et al., 2020; Haigo et al., 2003; Martin and Goldstein, 2014; Nikolopoulou et al., 2017), and severe defects in this process could result in an apparent increase in tissue width. We therefore used our previously reported *in toto* confocal imaging and computational segmentation approaches (Brooks et al., 2020) to examine cell apical areas (Figure 4A-C). These analyses revealed no difference in the average apical cell area (Figure 4D) or the distribution of apical cell areas (Figure 4E) within the lateral constrictive domains between control and *Lrp6* embryos. This normal level of apical constriction is also consistent with our observations of robust neural fold elevation in these mutants (Figure 1B, yellow arrowheads). Alternatively, failures in pseudo-stratification and the apicobasal thickening of the tissue are known to increase tissue width and block closure in some mutants (Grego-Bessa et al., 2016), but tissue thickness was unchanged in *Lrp6* embryos (Figure 4F). Thus, loss of *Lrp6* and the resulting impairment of Wnt signaling does not alter the patterned cell shape remodeling programs driving neural fold elevation, indicating that cranial closure defects arise from failures in other processes.

**Figure 4:**
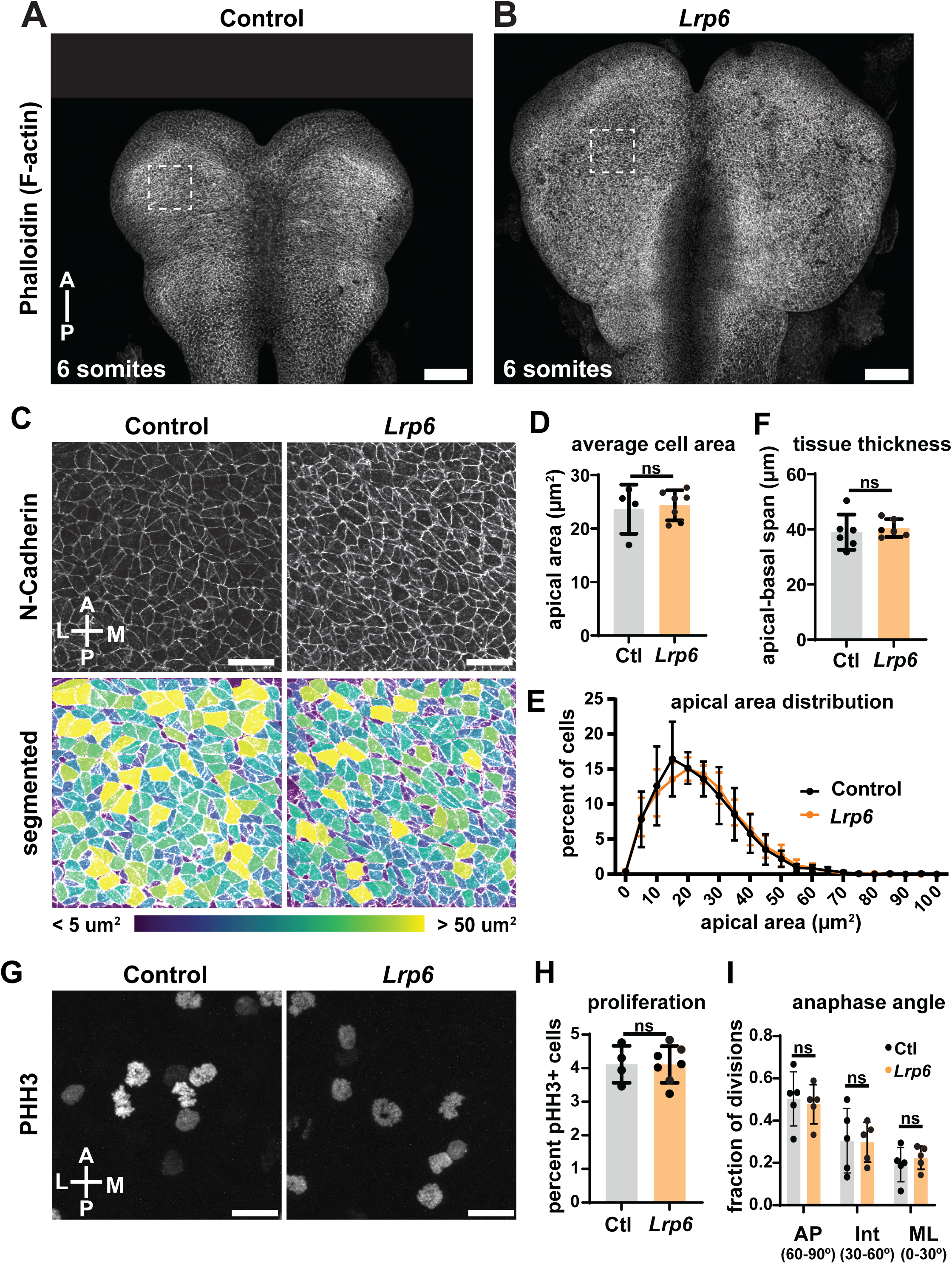
*Lrp6* mutants do not show defects in apical constriction, apicobasal elongation, or proliferation during elevation stages. (**A-B**) Maximum intensity projections of tiled confocal images of the future midbrain and hindbrain regions from control and *Lrp6* mutants at cranial neural fold elevation stages (E8.5, 5-7 somites). The apparent lengthening of the tissue in the *Lrp6* embryo is an artifact caused by the altered formation of the cranial flexure between the forebrain and midbrain tissues. (**C**) The apical areas of cells within the 100 µm x 100 µm boxed regions in A-B are shown by staining for the apical junctional marker N-Cadherin (top) and computational segmentation (bottom). (**D**) The average apical area of cells does not differ between control and *Lrp6* mutant embryos (control, 23.63 ± 4.59 µm^2^; *Lrp6*: 24.35 ± 2.81 µm. n = 4 control and 7 *Lrp6* embryos, p = 0.7492 by unpaired t-test, each dot represents the average of cell areas for a single embryo). (**E**) The distribution of cell apical areas does not differ between control and *Lrp6* mutants. (**F**) The apicobasal thickness of the lateral cranial tissues does not differ between control and *Lrp6* mutant embryos (control, 38.96 ± 6.40 µm; *Lrp6*: 40.44 ± 3.22 µm. n = 6 control and 6 *Lrp6* embryos, p = 0.6242 by unpaired t-test). (**G**) Staining for phospho-histone H3 (PHH3) in a 100 µm x 100 µm region of the lateral midbrain are shown. (**H**) No difference in the proportion of PHH3+ cells was observed between control and *Lrp6* mutants (control, 4.12 ± 0.55 percent; *Lrp6*: 4.11 ± 0.55 percent. n = 4 control and 7 *Lrp6* embryos, p = 0.9855 by unpaired t-test). (**I**) No difference in the orientation of anaphase separation angles was observed between control and *Lrp6* mutants by two-way ANOVA analysis with Sidak’s multiple comparisons. See Table S1 for n and p values. Anterior is up in all images, scale bars represent 100 µm in A-B, and 20 µm in C and G. ns, not significant; A, anterior; P, posterior; L, lateral; M, medial.

### The mammalian cranial neural plate is planar polarized, but this polarity does not depend on *Lrp6*

Convergent extension, driven by planar polarized cell intercalation, is a key driver of tissue narrowing that often depends on the Wnt-responsive planar cell polarity (PCP) pathway (Butler and Wallingford, 2017; Nikolopoulou et al., 2017; Sutherland et al., 2020). Defects in this process could theoretically explain the observed increase in tissue width in *Lrp6* mutants, but the relative strength of planar polarization in the mammalian cranial neural folds has not been extensively characterized, confounding analysis of this possibility. To address this gap, we developed an automated approach based on ridge detection algorithms ((Steger, 1998); see materials and methods) to quantitatively examine polarization patterns in the cranial neural folds. We first verified the utility of this method by examining the polarization of phospho-Myosin2 Regulatory Light Chain (pMRLC), a marker for the actively contractile myosin population that remodels cell junctions during cell intercalation. Consistent with our prior manual analysis (Brooks et al., 2020), automated detection showed moderate but highly reproducible polarization, with ∼40% of pMRLC enriched junctions and multicellular cables oriented within 30° of the mediolateral axis compared with only 25% oriented within 30° of the anteroposteior axis (Figure 5A-B). These cables ranged in length from ∼5 to ∼20 µm, and we wondered if shorter cables, representing ∼1-2 cell diameters, would show a more random orientation given they represented individual or paired cell edges. However, when separated for analysis both these short cables (≤10 µm) and longer multicellular cables (>10 µm) showed similar levels of polarization (Supplemental Figure 2A-B). We next expanded this analysis to the core PCP component VANGL2, which also exhibited moderate and length-independent planar polarization of multicellular cables (Figure 5D-E, Supplemental Figure 2C-D). Because our automated analysis only captured those cell edges and cables that exhibited enriched pMRLC and VANGL2, we next used a manual approach to analyze intensity along all edges within a 50 µm^2^ region of the lateral constrictive domain, using either phallodin or N- cadherin to identify junctions. In this analysis, edges aligned along the mediolateral axis (0-30°) showed 1.75-fold enrichment of pMRLC (Figure 5C) and 1.4-fold enrichment of VANGL2 (Figure 5F) over anteroposterior aligned edges (60-90°). Together with a previous analysis of the PCP component DVL2 (McGreevy et al., 2015), these data argue that the mammalian cranial neural plate shows significant and reproducible levels of planar polarization.

**Figure 5:**
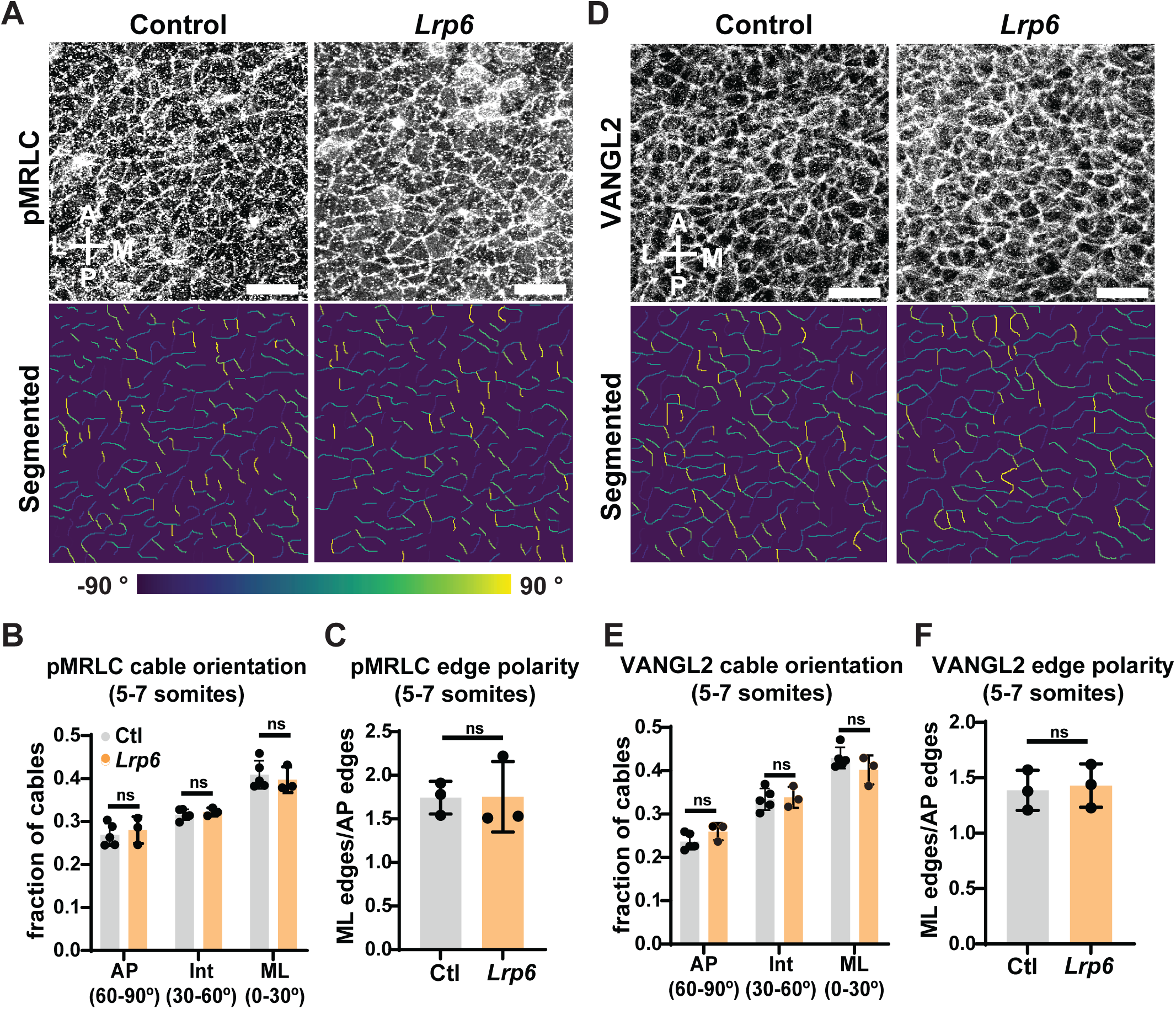
Planar polarization of cranial neural tissues is normal in *Lrp6* mutants. (**A**) Staining for phosphorylated myosin regulatory light chain (pMRLC, top) and segmented cables that are color coded according to angular orientation (bottom; −90° is posterior, 90° is anterior). Images are 100 µm x 100 µm lateral regions, similar to those analyzed in Figure 4. (**B**) The fraction of pMRLC cables in three angular bins, after taking the absolute value of the individual angles, is plotted for control and *Lrp6* mutants. Note that both control and *Lrp6* mutants show an equivalent bias toward mediolaterally aligned cables (0-30°, Ctl: 0.41; *Lrp6*: 0.40) over intermediate (30-60°, Ctl: 0.31; *Lrp6*: 0.32) and anteroposterior (60-90°, Ctl: 0.27; *Lrp6* 0.28) alignment. (**C**) Manual analysis of pMRLC cell edge intensities demonstrates that accumulation of pMRLC is also polarized, with mediolaterally aligned edges in both control and *Lrp6* mutant embryos exhibiting approximately 75% more pMRLC intensity on average than anteroposteriorly aligned edges (Ctl, 1.74 ± 0.19 fold enrichment; *Lrp6*, 1.75 ± 0.40; p = 0.9709 by unpaired t-test, n = 3 control, 3 *Lrp6* embryos). (**D**) Staining for the planar cell polarity component VANGL2 is shown in 100 µm x 100 µm lateral regions (top), and segmented cable orientation (bottom). (**E**) The fraction of VANGL2 cables in three angular bins, after taking the absolute value of the individual angles, is plotted for control and *Lrp6* mutants. Note that both control and *Lrp6* mutants show an equivalent bias toward mediolaterally aligned cables (0-30°, Ctl: 0.43; *Lrp6*: 0.40) at frequencies similar to that observed for pMRLC staining and biased over intermediate (30-60°, Ctl: 0.33; *Lrp6*: 0.34) and anteroposteriorly (60-90°, Ctl: 0.24; *Lrp6* 0.26) aligned ones. (**F**) Manual analysis of VANGL2 cell edge intensities demonstrates mediolaterally biased accumulation in both control and *Lrp6* mutant embryos (Ctl, 1.39 ± 0.18 fold enrichment; *Lrp6*, 1.43 ± 0.20; p = 0.7915 by unpaired t-test, n = 3 control, 3 *Lrp6* embryos). See Table S1 for n and p values. Anterior is up in all images, scale bars represent 20 µm. ns, not significant; A, anterior; P, posterior; L, lateral; M, medial.

We next tested whether loss of Wnt singling would impair tissue polarization, potentially explaining the increase in cranial neural fold width. However, neither pMRLC nor VANGL2 cable polarization decreased in *Lrp6* mutants, either across all cables (Figure 5A-B) or when separating short and long cables (Supplemental Figure 2A-D). Additionally, there was no significant difference in the number of detected cables for either staining (Supplemental Figure 2E-F) or change in mediolateral enrichment of either pMRLC or VANGL2 (Figure 5C, F). Together, these data indicate that *Lrp6* function is not required for planar polarization of the cranial neural plate. This surprising result may reflect the redundancy between LRP5 and LRP6 during Wnt signaling (Joeng et al., 2011; Kelly et al., 2004), an interpretation supported by the perdurance of some Wnt reporter activity in *Lrp6* mutants (Figure 2). Nonetheless, these analyses demonstrate that defects in planar polarization cannot explain the increased cranial tissue width or closure defects of *Lrp6* mutant embryos.

### An early increase in cell proliferation rates in early anterior neural tissues leads to cranial fold hyperplasia and terminal closure defects in *Lrp6* mutants

We next wondered if excessive cell growth or proliferation could explain the increased scale of the cranial neural tissues in *Lrp6* mutants. As discussed above, apical areas and cell heights do not differ in *Lrp6* mutant cranial tissues (Figure 4A-F), ruling out hypertrophy and prompting us to investigate proliferation and potential hyperplasia within the tissue. Consistent with the larger size of the tissue, *Lrp6* mutants showed a significant increase in the number of mitotically active cells within the future midbrain in absolute terms at elevation stages (Supplemental Figure 3A-C). However, when normalized to the total number of cells in the domain, these embryos showed an equivalent proliferation rate to controls (Figure 4G-H). The lack of increase in *relative* proliferation at these stages indicates that these excessive cell divisions are not the originating cause of the tissue scaling defect but rather a direct consequence of mis-scaling.

Alternatively, changes to the orientation of cell division have been proposed to underlie mediolateral widening of the tissue (Morriss-Kay, 1981) and Wnt signaling influences the orientation of cell division in several contexts (Di Bella et al., 2021; Kim et al., 2013; Schlesinger et al., 1999). We therefore tested whether the pattern of division orientation was changed by loss of *Lrp6*. In controls, mitotic separation events showed a significant bias toward anteroposterior alignment supporting models proposing anisotropic growth along this axis (Alvarez and Schoenwolf, 1991; Jacobson and Tam, 1982; Morriss-Kay, 1981). However, *Lrp6* mutants maintained this orientation bias (Figure 4I) while exhibiting a similar number of anaphase events (Supplemental Figure 3D) indicating that altered division orientation is not the cause of tissue widening.

Given that the known cellular drivers of cranial remodeling and neural fold elevation were all normal, and that the rate of proliferation was also normal at elevation stages, we reasoned that the terminal closure phenotype was likely to be a consequence of defects at earlier stages. This was supported by our observations that elevation stage specific loss of *Lrp6* did not prevent cranial closure (Figure 1C), and that cranial neural plate size was already significantly increased at pre-elevation stages (Figure 3A-B). We therefore investigated two key processes required to appropriately set the initial scale of the neural plate, cranial neural induction and early proliferation.

In the mouse, cranial neural tissues are induced by signals originating from the anterior visceral endoderm (AVE), which include inhibitors that protect early cranial tissues from the posteriorizing activity of Wnt signals (Levine and Brivanlou, 2007; Ozair et al., 2013; Stower and Srinivas, 2014). We reasoned that the reduction in Wnt signaling in *Lrp6* mutants might therefore lead to excessive anteriorization of the neural tissues at the expense of more posterior fates. Because the anterior-most tissues are significantly wider than more posterior regions, such a mis-fating may also lead to tissue scaling defects. To test this hypothesis, we used transverse sectioning to examine the distribution of OTX2, a key anterior fate determinant (Acampora et al., 1995; Ang et al., 1996; Matsuo et al., 1995; Rhinn et al., 1998), after the completion of induction at E7.5 (Figure 6A). In control embryos OTX2 occupied approximately 60% of the diameter of the embryonic cylinder, and we observed no change in proportional occupancy in *Lrp6* mutants (Figure 6C). We also examined the organization of β-catenin in OTX2+ anterior tissues at these stages and found that it was largely junctional and showed no significant differences between control and *Lrp6* mutant embryos (Supplemental Figure 4D-E). This is in line with a prior study of Wnt signaling dynamics in these tissues (Hernández-Martínez et al., 2019) and likely reflects the strong negative regulation of the pathway in this region. Together these analyses demonstrate that gross mis-patterning of anterior neural fates is not the cause of the tissue-scaling defects in *Lrp6* mutants.

**Figure 6:**
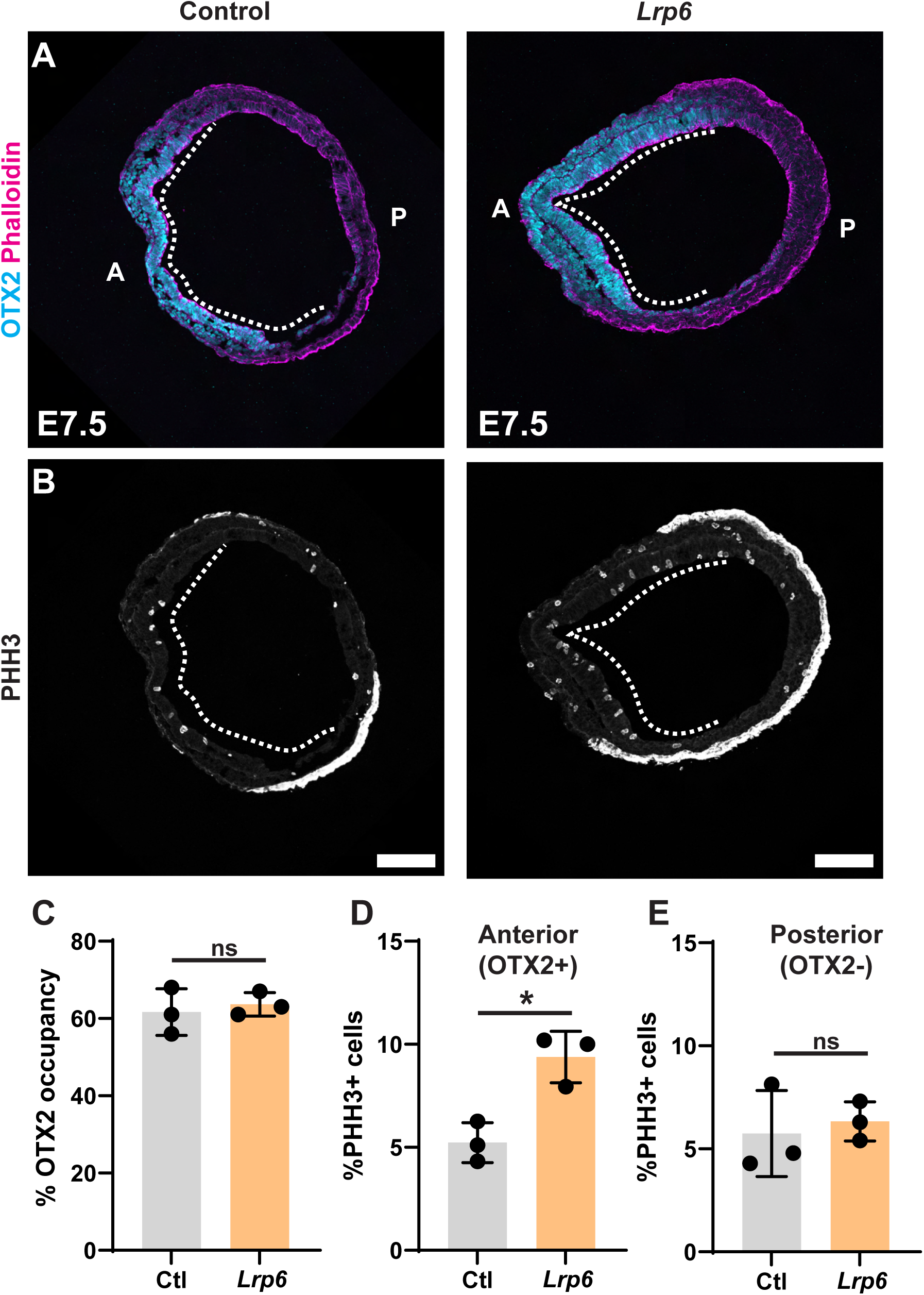
*Lrp6* mutants show excessive proliferation in anterior neural tissues at early neural plate stages. (**A-B**) Transverse sections through control and *Lrp6* mutant embryos at E7.5, stained with (A) an antibody against OTX2 and counterstained with Phalloidin (F- actin) and, or (B) the S-M phase marker phospho-HistoneH3. Anterior is to the left, dashed white lines indicate the OTX2+ neuroepithelial region. (**C**) Quantification of the percentage of the epiblast tissue marked with OTX2 shows that anterior fates are not expanded in *Lrp6* mutants (control: 61.67 ± 6.03 %; *Lrp6*: 63.67 ± 3.06 %; n = 3 control, 3 *Lrp6* embryos at E7.5; p = 0.6352 by unpaired t-test). (**D**) *Lrp6* mutants show a significant increase in the proportion of proliferative cells in the anterior (OTX2+) neural tissues as compared to controls (control: 5.23 ± 0.97 %; *Lpr6*: 9.38 ± 1.25%; n = 3 control, 3 *Lrp6* embryos; p = 0.0105 by unpaired t-test). (**E**) No difference is observed in the proportion of proliferative cells between control and *Lrp6* mutants in the posterior (OTX2-) neural tissues (control: 5.74 ± 2.08 %; *Lrp6*: 6.33 ± 0.95 %; n = 3 control, 3 *Lrp6* embryos; p = 0.6783 by unpaired t-test). Scale bars represent 100 µm; A, anterior; P, posterior.

Next, we returned to the question of excessive cell proliferation. Despite the data above demonstrating equivalent rates of proliferation between control and *Lrp6* mutants at later stages, an early burst of excessive proliferation could result in early hyperplasia that would then be maintained and expanded by normal ongoing proliferation in the tissue. To test this hypothesis, we examined the proportion of neural plate cells exhibiting phospho-HistoneH3 signatures after tissue induction at E7.5. Strikingly, we observed a near doubling of the anterior proliferative index in *Lrp6* mutants (Figure 6B, D). Increased proliferative capacity was unique to the anterior neural tissues, as the posterior proliferative index remained equivalent to controls (Figure 6B, E), and the underlying anterior mesoderm and the anterior visceral endoderm of *Lrp6* mutants did not exhibit statistically significant increases in cell proliferation rate (Supplemental Figure 4A-C). This increased proliferative capacity specifically in the cranial neural plate, integrated over the full tissue, is expected to result in the addition of a significant number of excessive cells, explaining the tissue hyperplasia observed at later stages. Further, this analysis suggests that the role of Wnt in tuning proliferation rates may be unique to anterior neural tissues, or that loss of Wnt may impair cell cycle control mechanisms specific to these tissues.

Altogether, our results indicate that loss of *Lrp6* leads to a temporally delimited increase in cell proliferation, which occurs specifically in early anterior neural tissues just after the completion of induction. This window of increased proliferation results in significant tissue hyperplasia prior to the onset of cranial neural fold elevation. Because the other cell dynamics of closure including apical constriction and planar polarization appear normal and neural fold elevation is robust, the likely genesis of the final cranial neural tube closure defect is an inability for these mechanisms to compensate for the increase is tissue width driven by early excessive proliferation. The increased tissue width, in turn, results in a failure of neural fold apposition over the midline, blocking fold fusion. Thus, these mutants reveal a previously unknown role for Wnt signaling in control of temporal patterns of cell proliferation as well as a novel pathogenic mechanism underlying cranial neural tube closure defects.

### Midbrain specific hyperactivation of Wnt signaling leads to altered actin organization and defective apical constriction resulting in neural fold elevation defects

In addition to the cranial closure defects observed upon reduced Wnt signaling, mutants that increase Wnt activity also prevent cranial closure (Gray et al., 2013; Kimura-Yoshida et al., 2015), but whether pathway reduction and hyperactivation result in equivalent defects at the level of patterned cellular dynamics is unknown. To address this question, we asked if elevating cranial tissues were sensitive to inappropriately increased levels of Wnt signaling. To do so we used a previously established mouse line (*Apc^tm1Tno^,* (Shibata et al., 1997)) capable of conditionally inactivating APC and thus hyperactivating Wnt signaling (Azzolin et al., 2014; Buchert et al., 2010; Martinez et al., 2009; Pawlikowski et al., 2013; Radulescu et al., 2013; Sansom et al., 2004; Shibata et al., 1997; Varnat et al., 2010). Because previous work indicates that *Apc* is broadly expressed throughout the epiblast (Ishikawa et al., 2003), we sought to use *Wnt1-Cre2* to drive APC truncation specifically in the midbrain at neural fold elevation stages in order to bypass earlier Wnt-dependent events. However, *Wnt1-Cre2* is also active in the male germline (Dinsmore et al., 2022) and a previous study showed hypomorphic effects in *Apc^tm1Tno^* genotypes consisting of one copy of the conditional allele and one copy of the truncated allele (Buchert et al., 2010). Therefore, to avoid these potentially confounding early hypomorphic effects, we only examined embryos from crosses where the Cre transgene was inherited from the dam.

Embryos with conditional APC inactivation specifically in elevating midbrain neural folds (*Apc^tm1Tno/tm1Tno^; Wnt1-Cre2*; hereafter called *Apc-cIN* for APC conditional inactivation) displayed highly penetrant exencephaly (9/13 mutants, ∼70%, Figure 7A). Of note, and in contrast to *Lrp6* mutants, *APC-cIN* mutants showed strong defects in neural fold elevation (compare Figure 7A vs 1B). This difference suggested that hyperactivation of Wnt signaling impairs cranial closure via a different cellular mechanism than that caused by reduced Wnt signaling. This hypothesis was further supported by the absence of cranial neural plate scaling defects in *APC-cIN* embryos (Figure 7B-C).

**Figure 7:**
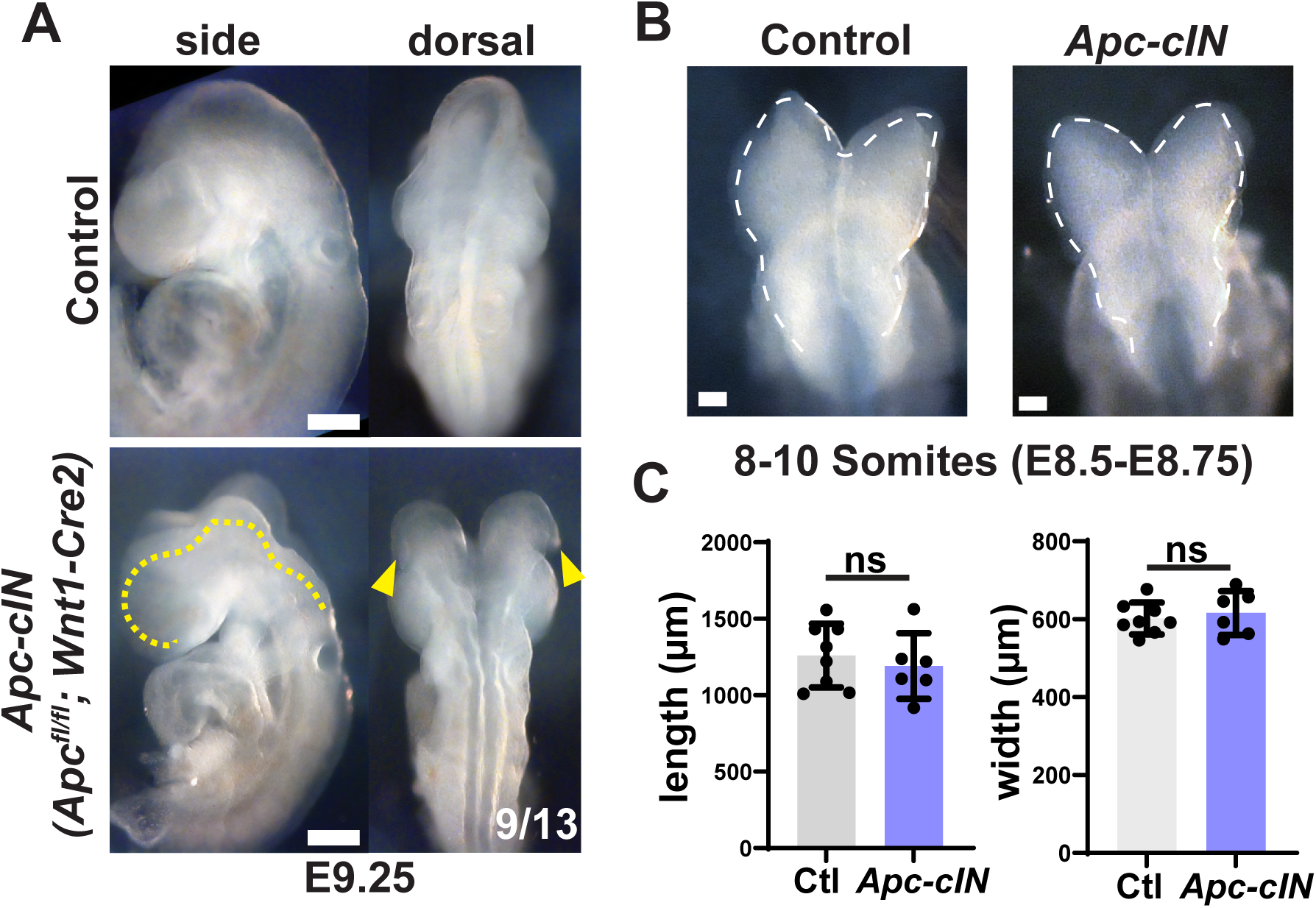
Cranial neural fold specific Wnt hyperactivation leads to neural fold elevation and cranial closure defects. (**A**) Conditional hyperactivation of Wnt signaling by forced truncation of APC specifically in the midbrain prevents cranial neural tube closure from the level of the forebrain to the hindbrain-spinal cord juncture (dashed yellow line), with high penetrance (∼70%). These *Apc-cIN* (short for APC conditional inactivation) mutants are distinct from *Lrp6* mutants due to strong defects in neural fold elevation (compare yellow arrowheads here and in Fig 1A). (**B**) Brightfield images of control and *Apc-cIN* embryos showing the mid- and hindbrain regions at 8 somites. (**C**) Hyperactivation of Wnt signaling by conditional APC truncation in *Apc-cIN* mutants does not lead to tissue scaling defects in either length (control, 1259 ± 208.6 µm; *Lrp6*: 1191 ± 214.7 µm. n = 8 control and 6 *Apc- cIN* embryos, p = 0.5604 by unpaired t-test) or width (control, 602.1 ± 41.09 µm; *Lrp6*: 616.2 ± 56.29 µm. n = 8 control and 6 *Apc-cIN* embryos, p = 0.5967 by unpaired t-test). Scale bars represent 100 µm, dashed outlines indicate edges of the neural plate. ns, not significant; **, p < 0.01, ***, p < 0.001. Anterior is up in all images.

We next wanted to assess the consequences of APC inactivation on Wnt signaling in this tissue. However, despite numerous crosses we never recovered embryos with both a copy of the Tcf/Lef reporter transgene and conditional APC truncation suggesting this combination may be genotoxic. As an alternative, we examined β-catenin organization given that APC inactivation disrupts its cytoplasmic destruction and nuclear export resulting in excessive Wnt target activation (Ha et al., 2004; Henderson, 2000; Munemitsu et al., 1995; Sansom et al., 2004; Staal et al., 2002; Su et al., 2008). A caveat to this analysis is that in some contexts APC has also been implicated in the regulation of β-catenin dependent cell-cell adhesion at junctions (Hamada and Bienz, 2002; Klingelhöfer et al., 2003). To begin differentiating between these roles for APC, we examined β- catenin organization at the level of apical junctions as well as at a more basal position within the tissue. At cell apices, junctional β-catenin showed a roughly two-fold enrichment over cytoplasmic signal in both control and *Apc-cIN* embryos (Figure 8A-B), suggesting that enrichment of β-catenin at the adherens junctions was not impaired by APC inactivation. More basally, *Apc-cIN* embryos exhibited an increase in the number of cells with obvious cytoplasmic β-catenin signatures and a general reduction of the ratio of junctional to cytoplasmic intensity compared to controls (Figure 8C-D). Notably, at this basal level of the tissue the vast majority of the cytoplasmic volume is occupied by the nucleus (Brooks et al., 2020; Grego-Bessa et al., 2016), and we saw no evidence for nuclear exclusion of β-catenin. Together, these data argue that the primary consequence of APC inactivation in this tissue is the accumulation of cytoplasmic and/or nuclear β-catenin, without obvious defects in junction organization or cell-cell adhesion.

**Figure 8:**
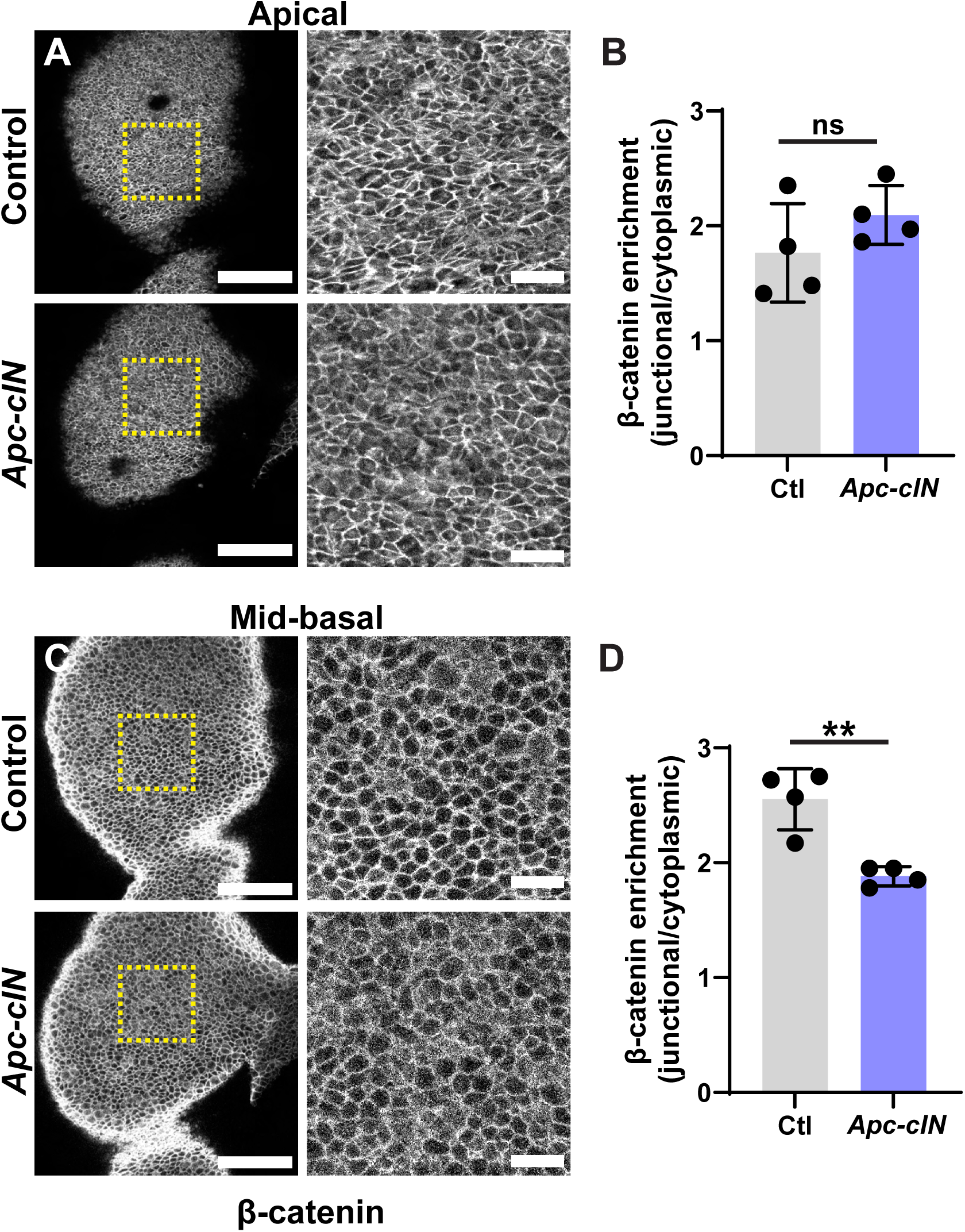
*Apc-cIN* mutants demonstrate increased cytoplasmic accumulation of β-catenin in basal domains at elevation stages. (**A**) The apical surface of the left midbrain cranial neural fold stained with an antibody against β-catenin is shown (left) along with a crop of the lateral constrictive region (right) for control and *Apc-cIN* embryos. (**B**) Quantification of the enrichment of junctional to cytoplasmic β-catenin at the apical surface reveals no significant difference between control and *Apc-cIN* embryos (control, 1.77 ± 0.43; *Apc-cIN*, 2.10 ± 0.26. n = 4 control and 4 *Apc-cIN* embryos, p = 0.2348 by unpaired t-test). (**C**) A single z-plane at the midpoint between the apical and basal surfaces of the left midbrain cranial neural fold stained with an antibody against β-catenin is shown (left) along with a crop (right) for control and *Apc-cIN* embryos. Note that many cells in the *Apc-cIN* embryo have apparent high β-catenin intensity in their cytoplasm. (**B**) Quantification of the enrichment of junctional to cytoplasmic β-catenin at this mid-basal position demonstrates a significant reduction in the junctional enrichment driven by the observed increase in cytoplasmic staining for the protein (control, 2.55 ± 0.27; *Apc-cIN*, 1.88 ± 0.08. n = 4 control and 4 *Apc-cIN* embryos, p = 0.0030 by unpaired t-test). Scale bars indicate 100 µm in parent images, 20 µm in crops. Anterior is up and medial is right in all images. ns, not significant; **, p < 0.01.

Because we and others have previously shown that cranial neural fold elevation requires patterned apical constriction (Brooks et al., 2020; Eom et al., 2011; Grego-Bessa et al., 2015; Kowalczyk et al., 2021; Lesko et al., 2021; Ohmura et al., 2012; Sulistomo et al., 2019), we next asked if *Apc-cIN* mutants displayed defects in this cell behavior. Indeed, these mutants demonstrated significant increases in average cell apical area and a shift in the distribution of areas towards larger apical domains (Figure 9A-G, Supplemental Figure 5A). These defects are equivalent in magnitude to those of mutants known to have neural fold elevation defects driven by failure of apical constriction (Brooks et al., 2020) and occurred in the absence of changes to cell height (Figure 9H) or changes in the relative proliferation rate between control and *Apc-cIN* mutants (Supplemental Figure 5B-C). Together, these data indicate that loss of APC function inhibits apical constriction, blocking cranial neural fold elevation and leading to terminal cranial closure defects.

**Figure 9:**
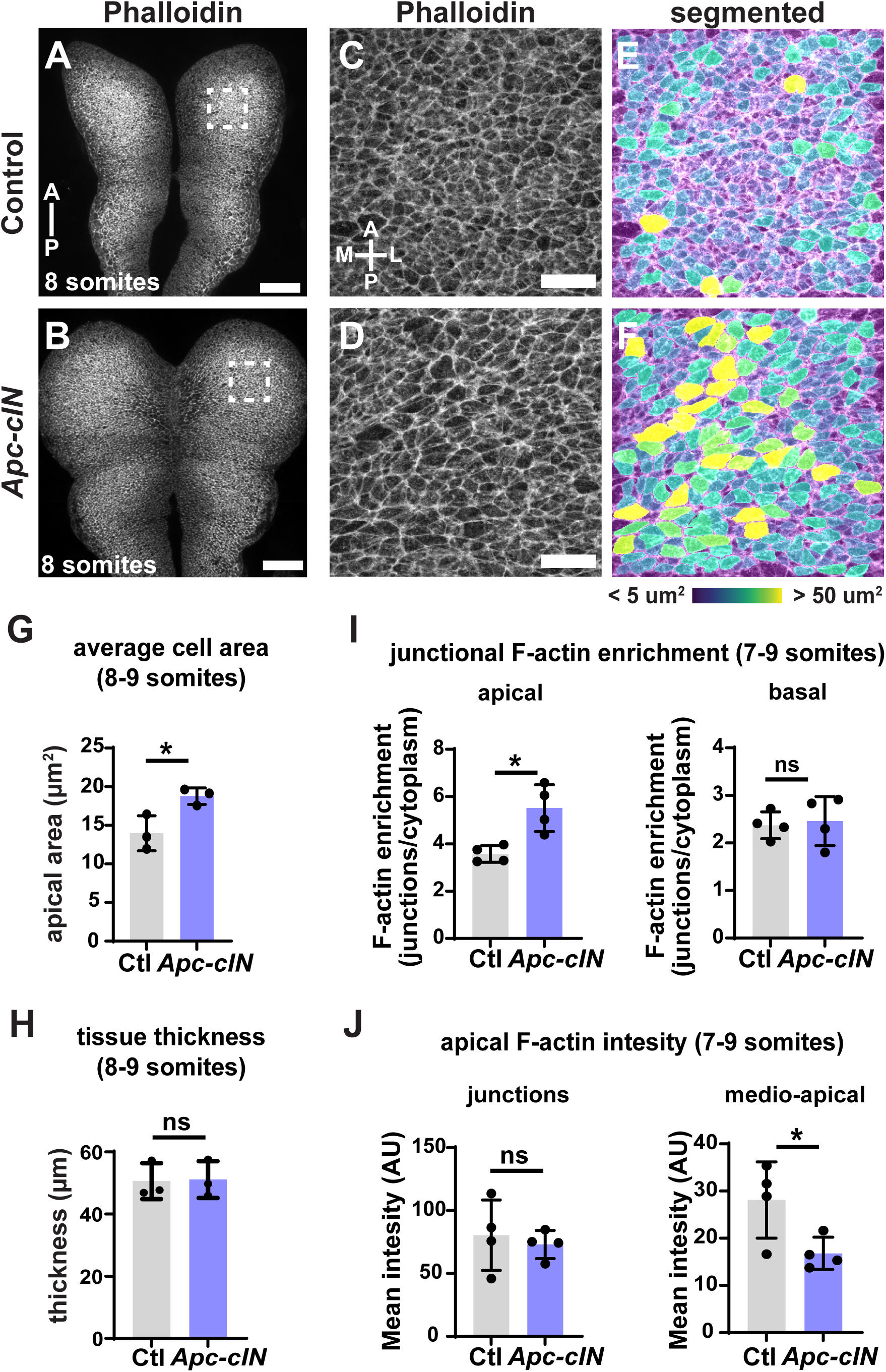
*Apc-cIN* mutants show failure of apical constriction and altered actin organization at elevation stages. (**A-B**) Maximum intensity projection of tiled confocal images of the midbrain and hindbrain regions of (A) control and (B) *Apc-cIN* mutant embryos at 8 somites. Dashed boxes indicate the areas analyzed in C and G. (**C-D**) The apical areas of cells within the 100 µm x 100 µm boxed regions in A-B are shown by staining for phallodin (F-actin). (**E-F**) Computational segmentation of apical areas shown in (C-D) (right). Note the increase in cells displaying very large apical areas in *Apc-cIN* mutants. (**G**) Quantification of the average apical cell area of embryos. *Apc-cIN* mutants show a significant enlargement of apical area (Ctl: 13.36 ± 2.29 µm^2^; *Apc-cIN*: 18.78 ± 1.07 µm^2^; n = 3 control and 3 *Apc-cIN* embryos; p = 0.0293). (**H**) There is no difference in neural plate thickness between control and *Apc-cIN* embryos (Ctl: 50.61 ± 5.77; *Apc-cIN*: 51.10 ± 5.92; n = 3 control, 3 *Apc-cIN*; p = 0.9238 by unpaired t-test). (**I**) *Apc-cIN* embryos show an increase in the junctional enrichment of F-actin specifically at the apical surface (Ctl: 3.57 ± 0.35 µm^2^; *Apc-cIN*: 5.52 ± 0.99; n = 4 control and 4 *Apc-cIN* embryos; p = 0.0100 by unpaired t-test), but basal actin organization does not differ in these mutants (Ctl: 2.37 ± 0.28; *Apc-cIN*: 2.46 ± 0.51; n = 4 control and 4 *Apc-cIN* embryos; p = 0.7765 by unpaired t-test). (**J**) The mean intensity of junctional F-actin (phalloidin) does not differ between control and *Apc-cIN* embryos (Ctl: 80.44 ± 27.96 arbitrary fluorescence units; *Apc-cIN*: 73.01 ± 11.22; n = 4 control and 4 *Apc-cIN* embryos; p = 0.6392 by unpaired t-test). However, the mean intensity of medio-apical cytoplasmic actin is significantly reduced in *Apc-cIN* mutants (Ctl: 28.10 ± 8.10 arbitrary fluorescence units; *Apc-cIN*: 16.80 ± 3.42; n = 4 control and 4 *Apc-cIN* embryos; p = 0.0422 by unpaired t-test).

We next sought to explore potential molecular underpinnings for these apical constriction defects. First, we examined the distribution of VANGL2 as—in addition to its role in planar polarization—it interacts with factors that control apical constriction (Kowalczyk et al., 2021). However, *Apc-cIN* embryos exhibited no changes in VANGL2 cable polarity, cable number, or mediolateral enrichment (Supplemental Figure 6A-D), indicating both that planar polarization is normal and that changes in VANGL2 cannot explain the apical constriction defects. Next, we explored potential defects in actomyosin contractility, which physically reshapes the apical surface area during constriction. Efficient constriction requires the regulated partitioning of F-actin between junctional and medio-apical cytoplasmic populations (Martin and Goldstein, 2014), and so we asked if this organization was altered in *Apc-cIN* embryos. We found that mutants demonstrated a robust increase in the ratio of junctional to cytoplasmic F-actin signal apically, but basal actin organization appeared to be normal (Figure 9I), indicating that actin organization is defective specifically within the constrictive apical domain. This apical imbalance was not associated with a change in actin planar polarization (Supplemental Figure 6E), leading us to directly examine the junctional and medio-apical actin populations, which revealed that junctional actin levels were similar between control and *Apc-cIN* embryos, but that mutants showed a significant reduction in medio-apical actin (Figure 9J). Given that this specific population is a critical regulator of apical constriction in many contexts, *e.g.* (Baldwin et al., 2023; Booth et al., 2014; Chung et al., 2017; Martin and Goldstein, 2014; Martin et al., 2009; Mason et al., 2013), we conclude that the failure to establish a robust medio-apical actin network is likely the cause of the apical constriction defects and resulting cranial closure failures of *Apc-cIN* mutants.

In sum, our data demonstrate that appropriate Wnt signaling levels are required for at least two distinct patterns of cell behavior required for cranial closure (Figure 9). First, there must be sufficient Wnt activity at post-induction stages to restrict proliferation in early anterior neural tissues. Impaired Wnt signaling, as in *Lrp6* mutants, leads to excessive cell proliferation at these stages and doubles the width of the tissue. The severe hyperplasia in these mutants prevents cranial closure despite the normal activity of other tissue remodeling mechanisms, including apical constriction and planar polarization. This tissue-scaling defect represents a novel cellular basis for cranial closure defects and may be specific to mutants that reduce Wnt signaling levels. Conversely, Wnt activity must be restricted at elevation stages, as hyperactivation of the pathway blocks apical constriction and prevents the tissue-wide curvature change required for neural fold elevation. Thus, Wnt levels must be correctly modulated to promote two key and temporally independent cell behaviors to reshape the cranial neural folds and drive closure.

## Discussion

Cranial neural tube closure is a critical event in vertebrate development, and failures in this process are among the most common and deleterious structural birth defects in humans (Wallingford et al., 2013; Zaganjor et al., 2016). Closure is driven by the active remodeling of the tissue by patterned cell behaviors, including shape change, rearrangement, and proliferation (Juriloff and Harris, 2018; Nikolopoulou et al., 2017). These patterns are strongly reliant on information provided by spatially restricted signals, such as morphogens, and failure in either these upstream signals or the downstream molecular regulators of cell activity lead to failed closure (Harris and Juriloff, 2007; Harris and Juriloff, 2010; Juriloff and Harris, 2018; Lee and Gleeson, 2020; Wilde et al., 2014). However, the precise consequence of perturbed morphogen signaling for downstream cell behaviors often remains opaque. Here, we showed that mutations resulting in either inappropriately reduced or hyperactivated Wnt signaling lead to defects in cranial closure. Intriguingly, defects in these two cases arose from different developmental trajectories. Loss of *Lrp6* led to an increase in cell proliferation specifically in the early cranial tissues, which led to significant hyperplasia at later stages. Because other known drivers of cranial closure including apical constriction, tissue thickening, and planar polarization were normal, and because *Lrp6* mutants showed strong cranial neural fold elevation, we conclude that the observed tissue hyperplasia creates a mechanical hindrance to final stages of closure that cannot be fully overcome by the normal function of these cell behaviors (Figure 10). This model is further supported by the lack of cranial closure defects upon conditional *Lrp6* ablation specifically at elevation stages, indicating that early defects in tissue size rather than direct disruption of elevation mechanisms underly the terminal closure phenotypes. Conversely, conditional hyperactivation of the pathway through controlled truncation of APC did not alter tissue scale but instead led to defects in actin organization which prevented apical constriction, leading to a failure of neural fold elevation and terminal cranial closure defects (Figure 10). Thus, Wnt signaling levels are critical determinants of two distinct events during closure, and pathway activity must be appropriately regulated throughout early cranial development to ensure both early tissue scaling, and later elevation occur appropriately.

**Figure 10:**
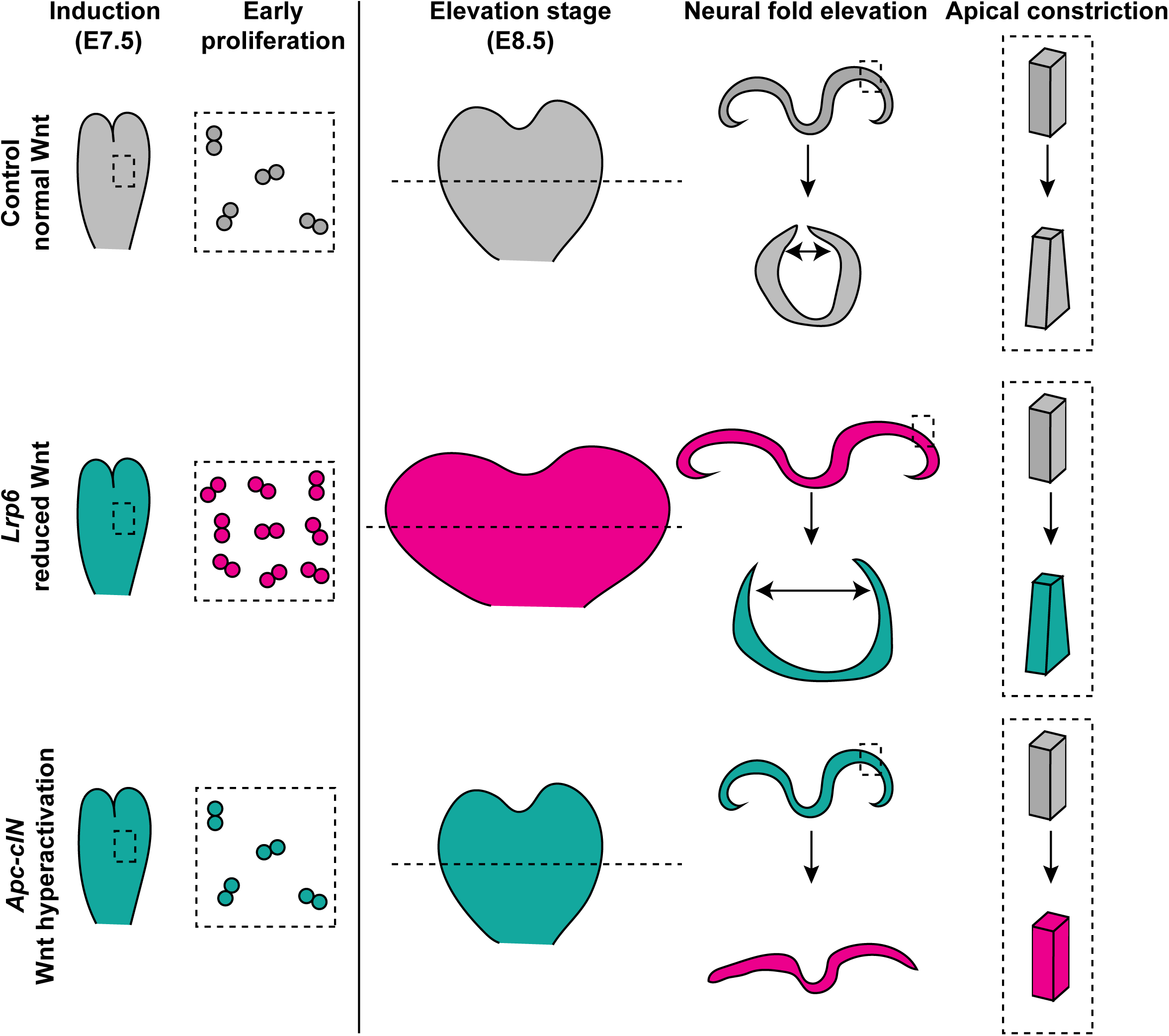
Model for the role of Wnt signaling levels in cranial neural tube closure. Schematic representation of the impact of Wnt reduction (*Lrp6* mutants) or hyperactivation (*Apc-cIN*) on elements of cranial neural tube closure. Events that are comparable with controls are highlighted in green, while those that are defective are highlighted in magenta. In *Lrp6* mutants, neural induction is normal, but early tissues show roughly double the rate of cell proliferation. This leads to a doubling of tissue width at elevation stages, and a resulting failure to bring the neural folds together over the midline despite the normal activity of apical constriction and neural fold elevation. Wnt hyperactivation in *Apc-cIN* mutants does not impact early events or tissues scale but blocks later apical constriction resulting in a failure of neural fold elevation. Thus, both conditions lead to cranial closure defects but do so via disruption of unique cellular mechanisms.

The fact that loss of LRP6 and concomitant reduction in Wnt signaling leads to excessive proliferation in early stages is noteworthy given that Wnt activity is generally pro-proliferative, including in neural tissues (Ikeya et al., 1997; Kalani et al., 2008; Reya and Clevers, 2005; Steinhart and Angers, 2018). In line with this, *Lrp6* mutants which show reduced Wnt signaling demonstrate a slight reduction in proliferation just after closure (Gray et al., 2013). Why then do early stages in these mutants show increased proliferative capacity? In one model, decreased Wnt signaling leads to an anteriorization of the tissue at a level downstream of the broad early patterning by OTX2, possibly by driving over expression or expansion of one of the anterior homeobox factors that control early neural patterning. Given that these homeobox factors can also influence proliferation rates (Andreazzoli et al., 2003; Gestri et al., 2005), this may explain the tissue hyperplasia of *Lrp6* mutants. In another model, reduced Wnt signaling could alter the timing or efficacy of the transition from epiblast to pro-neural fates, perhaps incorrectly maintaining a more pluripotent and proliferative state. Finally, studies in other developing tissues and during tumorigenesis have shown that there is significant cross talk between the Wnt and Hippo signaling pathways (Azzolin et al., 2012; Azzolin et al., 2014; Doddihal et al., 2024; Heallen et al., 2011; Varelas et al., 2010), and the proliferation and tissue scaling defects in *Lrp6* could result from secondary impacts on Hippo pathway activity. Regardless of the molecular basis for hyperproliferation, our data argue that proliferation is actively modulated in cranial tissues throughout closure stages. However, it remains unclear whether cell division *per se* is an active driver of their morphogenesis. Experiments directly increasing or decreasing proliferative capacity will help delineate the role(s) of cell division during closure.

To our knowledge, *Lrp6* mutants exhibit the earliest developmental onset for cranial closure defects reported to date, and the first directly attributable to altered cell proliferation rates at these early stages. This includes our own observations of mutants with increased Sonic Hedgehog signaling (Brooks et al., 2020), which do not show altered proliferation rates despite this pathway being generally pro-mitogenic (Corrales et al., 2004; Fink et al., 2018; Ishibashi and McMahon, 2002). Mutants that inhibit apoptosis, e.g. *Jnk1/2* doubles (Kuan et al., 1999; Sabapathy et al., 1999), may result in similar tissue scaling defects, as defects in this process result in neural tube hyperplasia at later stages (Cecconi et al., 2008), but the stage of hyperplasia onset in these mutants has not been identified. Another potentially interesting comparison comes from studies of hemifacial hypertrophy, a rare human developmental disease characterized by unilateral overgrowth of facial tissues (Dattani and Heggie, 2021). This disease has been proposed to stem from an increase in the number of neural crest cells due to unilateral hyperplasia of a single cranial neural fold (Pollock et al., 1985), though it remains unknown if this is a unifying etiology for the disease or if Wnt signaling is meaningfully altered in these patients.

Our investigation also found that Wnt hyperactivation leads to apical constriction defects. While our model is that these apical constriction defects arise from altered Wnt target expression, the observed defects could also potentially arise from a more direct modulation of junctional dynamics by APC (Bienz and Hamada, 2004; Hamada and Bienz, 2002; Klingelhöfer et al., 2003). We favor the transcriptional model due to our finding that β-catenin accumulated in cytoplasmic/nuclear compartments of *Apc-cIN* embryos, but junctional organization of the protein appeared normal. Additionally, hypermorphic *Lrp6* mutants—which are unlikely to directly inhibit any role of APC in junctional β-catenin organization—show defects in actomyosin organization in the neural plate (Allache et al., 2014; Gray et al., 2013) that are likely to result in failures of apical constriction comparable to our observations in *Apc-cIN* embryos.

Our model also suggests that the cell-level consequences of Wnt hyperactivation are very similar to those observed in mutants with excessive Shh signaling (Brooks et al., 2020). This is surprising given that Wnt and Shh are antagonistic in neural patterning (Kicheva and Briscoe, 2023; Ulloa and Martí, 2010), and we and others have recently shown that Shh activation inhibits transcription of several Wnt pathway members in neural tissues at closure stages (Brooks et al., 2024; Kim et al., 2019). In fact, based on these observations one of our initial hypotheses was that Wnt would act as a positive signal for apical constriction in opposition to Shh as a negative signal. However, our data instead indicate that high levels of activity in either pathway inhibit constriction, suggesting that each likely acts to set cell morphology in its own unique domain—the lateral edge of the tissue for Wnt and the midline for Shh—but must be restricted in the intervening domain. This model is further supported by our observation that reduced Wnt signaling in *Lrp6* mutants did not result in defects in apical constriction. An important caveat to this interpretation is that loss of *Lrp6* alone is not sufficient to completely abrogate Wnt signaling in these tissues, and studies of stronger loss of function mutants will be required to fully define if basal pathway activity is required for constriction. In either case, it remains unclear whether Wnt and Shh signals converge on shared downstream targets or instead control unique morphogenetic cassettes. Direct comparison of the actomyosin organization and gene-expression consequences of perturbations to these pathways will be required to address this important open question.

Another surprising result from this work is the lack of any obvious planar polarization phenotypes in the cranial neural plate of either *Lrp6* or *Apc-cIN* mutants, especially given that previous studies have found that LRP6 plays a role in convergent extension during gastrulation (Tahinci et al., 2007) and its loss can exacerbate spinal neural tube defects in *Vangl2* heterozygotes (Allache et al., 2014). Further, *Snx3* mutants have impaired Wnt signaling and exhibit defects consistent with convergent extension (Brown et al., 2020). All together these studies suggest that Wnt levels are important regulators of planar polarity in multiple tissues, including the posterior neural plate. However, we found that neither reduction nor hyperactivation of the Wnt pathway resulted in defective planar polarization of cells within the cranial neural tissues. Exciting recent work has demonstrated that the cranial and spinal neural plate have fundamentally different dynamic properties despite being continuous with one another (Baldwin et al., 2022; Christodoulou and Skourides, 2022), and perhaps the requirement for Wnt in planar polarization also differs along this axis. However, the caveat that *Lrp6* mutants do not fully ablate pathway activity in this tissue applies here as well, and studies using stronger loss of function approaches will be required to more fully address this question. Such studies will also be useful in determining the relative contribution of polarized cell intercalation to cranial morphogenesis, a critical gap in our current knowledge.

To conclude, this study lays the groundwork for several avenues of further investigation. First, it will be important to understand if and how Wnt signaling levels change in space and time as the tissue proceeds from induction to closure to distinguish between models for how proliferation is controlled. Second, examination of the intersection between Shh and Wnt signaling may reveal the molecular logic that governs the unique pattern of apical constriction observed in cranial tissues. Finally, further investigation of the Wnt pathway may provide insights into how tissues are appropriately scaled during development. This is a critical concern, as the cellular mechanisms driving morphogenesis must be accurately scaled to underlying tissue dimensions in order to appropriately reshape tissues for function during embryogenesis but the mechanisms controlling such scaling are not fully understood.

## Materials and Methods

### Mouse handling

Mice were bred and housed in accordance with the Guide for the Care and Use of Laboratory Animals of the National Institutes of Health and an approved Institutional Animal Care and Use Committee protocol (22-124) of North Carolina State University.

Timed pregnant mice were euthanized at E7.5-E10.5. Noon on the day of the vaginal plug was considered E0.5, and individual embryos were staged by counting the number of somites, or by Thieler criteria in stages prior to the appearance of somites. Strains used in this study included the following previously reported lines: the conditional *Lrp6^fl^* line [Lrp6^tm1.1Vari^, MGI ID: 5299216] (Joeng et al., 2011); the *Apc^tm1Tno^* line [Apc^tm1Tno^, MG ID: 1857966] (Shibata et al., 1997); the Tcf/Lef H2B-GFP transgenic insertion [Tg(TCF/Lef1-HIST1H2BB/EGFP)61Hadj/J, MGI ID: 4881506] (Ferrer-Vaquer et al., 2010); and the *Wnt1-Cre2* transgenic insertion line [E2f1^Tg(Wnt1-cre)2Sor^, MGI ID: 5485027] (Lewis et al., 2013). Genotyping was performed in accordance with reported protocols for each strain. Mice were received and maintained on a mixed background. Control embryos were wild-type and heterozygote littermates for global *Lrp6* analyses, or wild-type, heterozygote, and Cre-negative littermates for *Lrp6; Wnt1-Cre2* and *Apc-cIN* analyses.

### Whole-mount immunostaining

Embryos were dissected in ice-cold phosphate buffered saline (PBS) and fixed in 4% paraformaldehyde for either 1 hour at room temperature or overnight at 4 °C. Embryos were then washed 3x for 30 minutes in PBS + 0.1% Triton X-100 (PBTriton) at room temperature, blocked for 1 hour at room temperature in PBS containing 10% bovine serum albumin and 0.1% Triton X- 100 (blocking solution), and incubated with primary antibodies in blocking solution overnight at 4 °C. Embryos were then washed 3x 30 minutes in PBTriton at room temperature, incubated with secondary antibodies in blocking solution for either 1h at room temperature or overnight at 4 °C, and finally washed 3x 30 minutes in PBTriton. Primary antibodies used in this study were: rabbit anti-N-cadherin (1:500, Cell Signaling #13116), mouse anti-phospho-HistoneH3 (1:500, Cell Signaling #9706), rabbit anti-phospho-Myosin Regulatory Light Chain 2 (1:100, Cell Signaling 3674), rat anti-VANGL2 (1:100, Millipore MABN750), and goat anti-OTX2 (1:500, R&D Systems AF1979). Alexa conjugated Phalloidin and DAPI (ThermoFisher) were used as counterstains.

### Cryosectioning

E7.5 Embryos were dissected and fixed in 4% paraformaldehyde for 1 hour at room temperature, washed 3x 30 minutes in PBTriton and then incubated in 15% sucrose overnight followed by another overnight incubation in 30% sucrose. Embryos were then placed in cryoblocks containing OCT (Tissue-Tek) then frozen on dry ice and stored at −80 °C until sectioning. Embryos were sectioned on a cryostat (Leica) in 10-14 µm increments and sections were adsorbed onto Superfrost slides (Fisher). Slides were washed 3x 5 minutes in PBTriton at room temperature, blocked for 15 minutes with blocking solution (see above), incubated with primary antibodies for 30 minutes, washed 3x 5 minutes with PBTriton, incubated with secondary antibodies for 30 minutes, and washed 3x 5 minutes in PBTriton. Stained sections were then mounted in Prolong Gold antifade reagent (ThermoFisher).

### Brightfield imaging and morphometric analysis

Embryos were placed in a petri dish with a layer of 2% agarose on the bottom and filled with PBTriton. Small gouges were made in the agarose to gently restrain embryos in the desired orientation, and images were taken using a Zeiss Lumar V12 Steroescope equipped with a CCD camera. Cranial neural plate morphometrics were measured using FIJI/ImageJ (Schindelin et al., 2012; Schneider et al., 2012). The length of the cranial tissues was measured as a segmented line along the neural plate border from the anterior neural ridge to the otic sulcus. The width of the tissue was measured by drawing a line between the lateral neural plate borders at the widest position in the future midbrain.

### Confocal Microscopy

For whole mount imaging, embryos were mounted dorsal side down on #1.5 coverglass in PBTriton in Attofluor chambers (ThermoFisher A7816) using a small fragment of broken cover glass with dabs of vacuum grease to gently hold the embryo against the glass. Embryos were imaged on a Zeiss LSM 980 confocal microscope equipped with a Plan-NeoFluar 40x/1.3 oil immersion objective. Images were captured using tile-based imaging to acquire contiguous z-stacks of 50-100 µm depth, with an optical slice thickness of 0.6 µm and z-steps of 0.3 µm. Tile stitching and maximum intensity projection were performed in Zeiss ZEN Blue software. Imaging of cryosections was performed on the same microscope using a 20x/0.8 air immersion objective, with z-stacks of 10-14 µm depth imaged with a 1.8 µm optical slice thickness and a 0.9 µm z-step.

### Image analysis and quantification

All quantifications were performed on unprocessed images. To quantify Wnt response, Tcf/Lef-H2B-GFP intensity was normalized to DAPI in a contiguous series of 20×100 µm boxes along the midbrain stripe of expression in maximum intensity projections. We were unable to directly analyze the impact of APC truncation on Wnt signaling levels as, despite numerous crosses, we never recovered embryos that had both homozygous APC truncation and a copy of the Tcf/Lef:H2B-GFP reporter. This could indicate that Wnt hyperactivation in combination with this reporter transgene is embryotoxic.

Apical cell area was quantified in 100 µm x 100 µm regions in maximum intensity projections in a pair of regions on either side of the midline, approximately halfway between the midline and lateral edge, and halfway between the pre-otic sulcus and the cranial flexure. Cells entirely contained within these regions were segmented using CellPose (Stringer et al., 2021). Cell areas were quantified and cell area maps were generated using the MorphoLibJ plugin (Legland et al., 2016) in FIJI/ImageJ (Schindelin et al., 2012; Schneider et al., 2012). Tissue thickness was measured in orthogonal reconstructions of z-stacks, midway between the midline and lateral edge of the tissue.

Planar polarization of pMRLC and VANGL2 cables was measured by first using a ridge detection algorithm (Steger, 1998) in FIJI/ImageJ to identify linear junctional signals in a 100 µm x 100 µm region, selected as above, of maximum intensity projections of these stainings. After segmentation, a fit ellipse was generated for each detected object. In this analysis 90° is anterior, −90° is posterior, and after taking the absolute value, reported values range from 0-90°. The frequency of objects falling into one of three angular bins (anteroposterior, 60-90°; intermediate 30-60°; and mediolateral 0-30°) was calculated. For manual analysis, pMRLC or VANGL2 intensities along all cell edges in a 50 μm^2^ region of the lateral cranial neural plate were manually measured using the line tool in FIJI/ImageJ. Intensities along all edges with angles of 0-30° (mediolateral) and 60-90° (anteroposterior) from one embryo were averaged and then mediolateral enrichment was quantified by dividing these two values.

For analysis of junctional enrichment of β-catenin and F-actin (phalloidin) at elevation stages (E8.5), the mean junctional signal of each cell in a 30 μm^2^ region of the lateral neural plate was measured using the segmented line tool in FIJI/ImageJ to highlight the perimeter of the cell. Then the mean intensity from a polygon enclosing the majority of the cytoplasmic domain but excluding the junctions was recorded. Cell-wise junction/cytoplasm enrichment was calculated and then all individual cell values were averaged to generate a single value per embryo. A similar approach was used on β-catenin staining in transverse sections of E7.5 embryos, except instead of a 30 μm^2^ region, 30 cells in the OTX2+ domain were analyzed per embryo.

For elevation stages, the total number of phosphor-HistoneH3 cells between the cranial flexure and the pre-otic sulcus in one lateral neural fold was measured. The proportion of proliferative cells was calculated by normalizing the number of phospho-Histone H3 positive cells to the total number of cells, marked by DAPI, within a 100 µm x 100 µm region selected as described above. Anaphase angle separation was quantified in DAPI staining and all obvious chromosomal separation events in one lateral half of the cranial neural plate were quantified. The frequency of objects falling into one of three angular bins (anteroposterior, 60-90°; intermediate 30-60°; and mediolateral 0-30°) was calculated. For pre-elevation stages, the relative percentage of the tissue showing OTX2 staining was calculated by drawing a segmented line in FIJI/ImageJ, and sections were subsequently divided into two regions, anterior (OTX+) and posterior (OTX2-) and the number of PHH3+ cells to the total cells, specifically within the neuroepithelial tissues, was calculated independently in these regions. The same approach was used to calculate the proliferative rate in the mesoderm and anterior visceral endoderm tissues.

### Statistics

Statistical analysis and graph generation were performed in Prism software (GraphPad). All results are reported as mean ± standard deviation. Summary significance levels are: ns, not significant; * p<0.05; ** p<0.01; *** p<0.001. Statistical tests were the unpaired students t-test for comparing two conditions, and two-way ANOVA with either Tukey’s multiple comparisons or Sidak’s multiple comparisons. Details on the statistical tests, n values, and p values for each experiment can be found in Table S1. No formal power analyses were conducted, an n of 3-6 embryos per genotype was targeted for each analysis.

## Figure Assembly

Display images in figures were linearly adjusted to increase brightness and contrast in either FIJI/ImageJ or Photoshop (Adobe). Figures were assembled in Illustrator (Adobe).

## Acknowledgements

*Lrp6*^fl^ mice were a kind gift of Bart Williams (Van Andel Institute). We thank Kristina Borys for technical assistance, Lilian Lamech and Nanette Nascone-Yoder for thoughtful comments on the initial manuscript, Gillian Welch for support during revision, and our reviewers, whose thoughtful comments significantly improved this study. This work was supported by startup funds to ERB from the Department of Molecular Biomedical Science and the College of Veterinary Medicine at North Carolina State University.

## Figure Legends

**Supplemental Figure 1:**
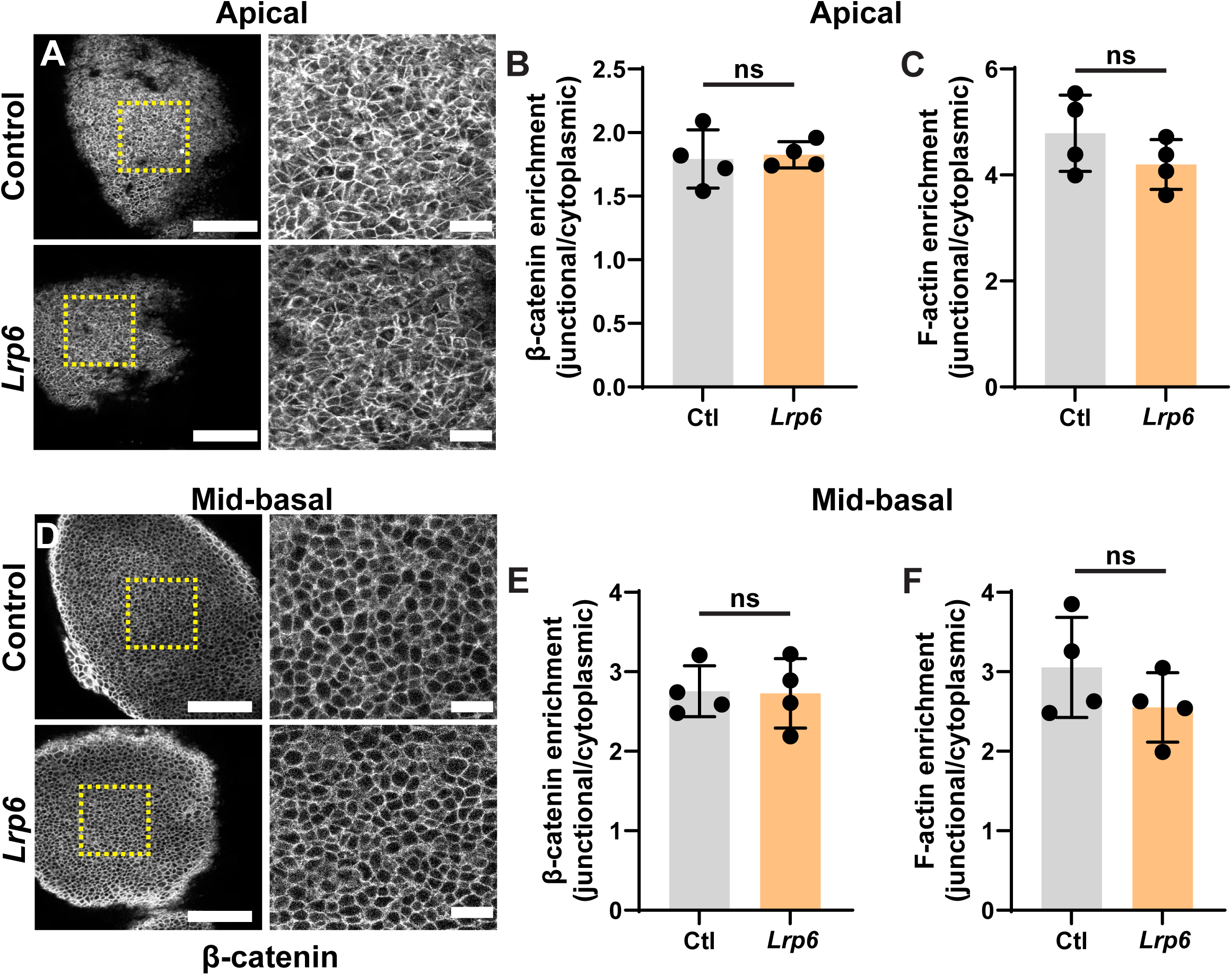
*Lrp6* mutants do not show changes to β-catenin or F-actin organization, or to the number of anaphase events, pMRLC cables, or VANGL2 cables observed in the midbrain neural folds. (**A**) The apical surface of the left midbrain cranial neural fold stained with an antibody against β-catenin is shown (left) along with a crop of the lateral constrictive region (right) for control and *Lrp6* mutant embryos. (**B**) Quantification of the enrichment of junctional to cytoplasmic β-catenin at the apical surface reveals no significant difference between control and *Lrp6* embryos (control, 1.79 ± 0.23; *Lrp6*, 1.82 ± 0.10. n = 4 control and 4 *Lrp6* embryos, p = 0.8048 by unpaired t-test). (**C**) The enrichment of F-actin at apical junctions compared to the apical cytoplasmic domain does not significantly differ between control and *Lrp6* mutant embryos (control, 4.79 ± 0.72; *Lrp6*, 4.20 ± 0.47. n = 4 control and 4 *Lrp6* embryos, p = 0.2185 by unpaired t-test). (**D**) A single z-plane at the midpoint between the apical and basal surfaces of the left midbrain cranial neural fold stained with an antibody against β-catenin is shown (left) along with a crop (right) for control and *Lrp6* embryos. (**E**) There are no significant differences in β-catenin organization between control and mutant embryos (control, 2.76 ± 0.32; *Lrp6*, 2.72 ± 0.08. n = 4 control and 4 *Lrp6* embryos, p = 0.9225 by unpaired t-test). (F) The enrichment of F-actin at mid-basal junctions compared to the equivalent cytoplasmic domain does not significantly differ between control and *Lrp6* mutant embryos (control, 3.06 ± 0.63; *Lrp6*, 2.55 ± 0.44. n = 4 control and 4 *Lrp6* embryos, p = 0.2369 by unpaired t-test) Scale bars indicate 100 µm in parent images, 20 µm in crops. Anterior is up and medial is right in all images.

**Supplemental Figure 2:**
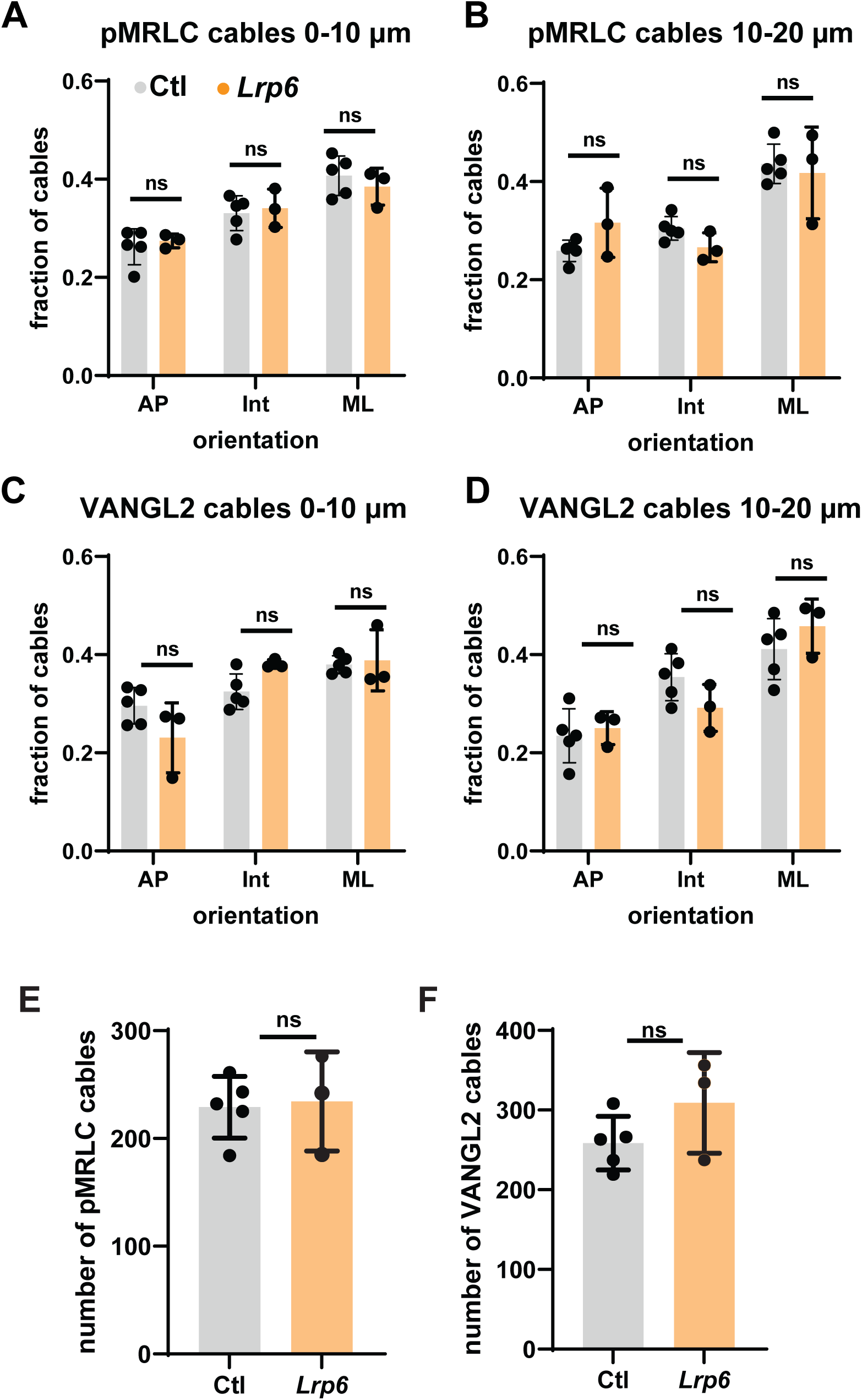
Polarity does not differ between either short or long pMRLC and VANGL 2 cables in *Lrp6* mutants. (**A-B**) The short (A, 0-10 µm) and long (B, 10-20 µm) cables of pMRLC identified by ridge detection are equivalently polarized in control and *Lrp6* mutants by two-way ANOVA with Sidak’s multiple comparisons. (**C-D**). The short (C, 0-10 µm) and long (D, 10-20 µm) cables of VANGL2 identified by ridge detection are equivalently polarized in control and *Lrp6* mutants by two-way ANOVA with Sidak’s multiple comparisons. See table S1 for bin means, n, and p-values. (**E**) Quantification of the number of pMRLC cables observed in a 100 µm x 100 µm region in control and *Lrp6* mutant embryos. (control: 229.0 ± 28.6; *Lrp6*: 234.3 ± 46.0; n = 5 control, 3 *Lrp6* embryos; p = 0.8432 by unpaired t-test). (**F**) Quantification of the number of VANGL2 cables observed in one 100 µm x 100 µm region in control and *Lrp6* mutant embryos. (control: 258.6 ± 33.7.6; *Lrp6*: 309.0 ± 63.3; n = 5 control, 3 *Lrp6* embryos; p = 0.1823 by unpaired t-test).

**Supplemental Figure 3:**
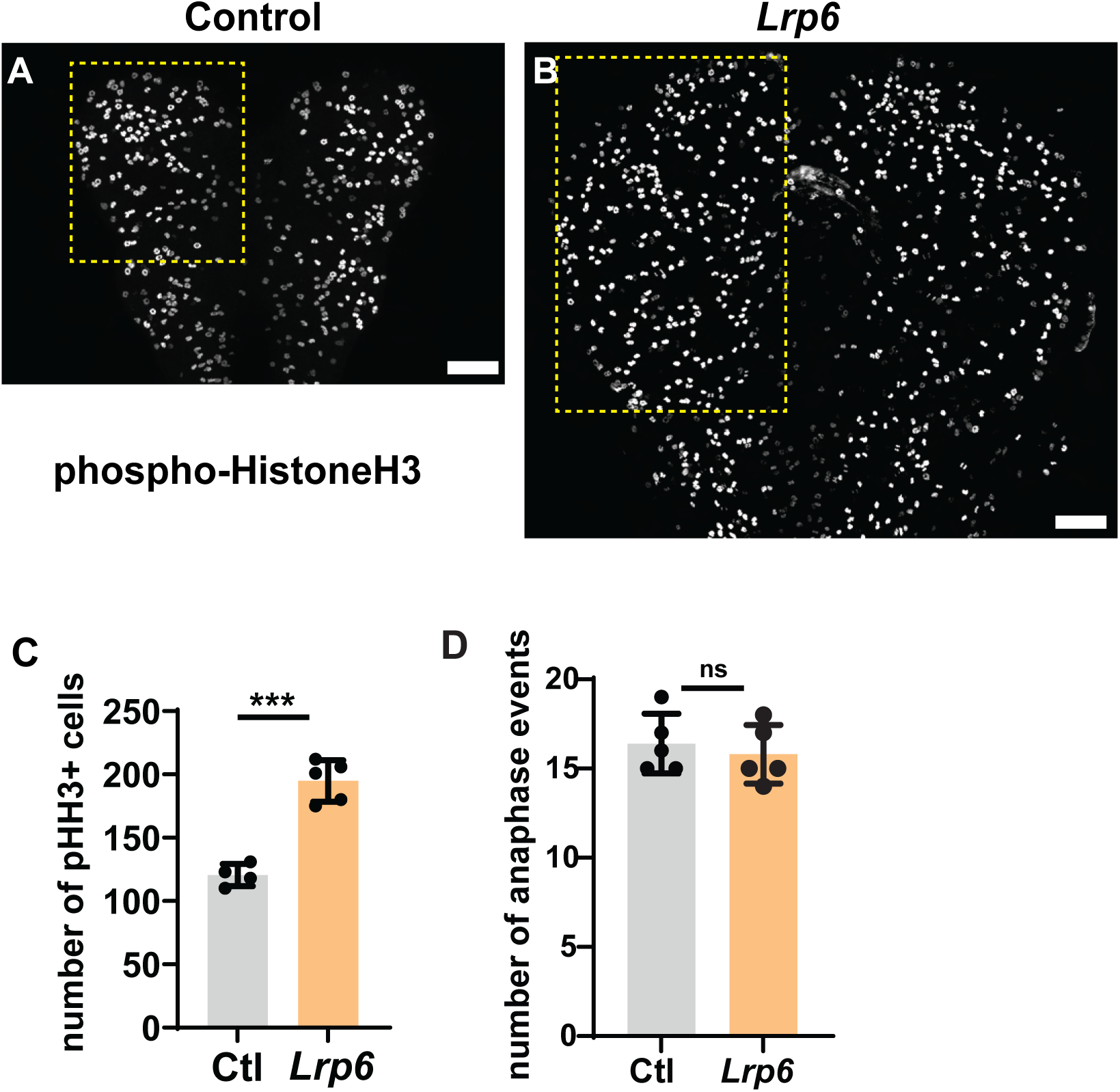
*Lrp6* mutants show an increase in the absolute number of mitotically active cells in the cranial neural plate at elevation stages. (**A-B**) Maximum intensity projections of cranial neural plates from control and *Lrp6* mutant embryos stained with the mitotic marker phospho-HistoneH 3. Dashed yellow boxes represent the midbrain regions used to quantify mitotic events. (**C**) Quantification of the total number of phospho-HistoneH3 positive cells in the future midbrain region of cranial neural plates in control and *Lrp6* mutant embryos (control, 120.5 ± 8.81 cells; *Lrp6*, 194.8 ± 16.36. n = 4 control and 5 *Lrp6* embryos, p < 0.0001 by unpaired t-test). (**D**) Quantification of the number of anaphase separation events observed in DAPI staining on one half of the cranial neural plate in control and *Lrp6* mutant embryos. (control: 16.4 ± 1.7; *Lrp6*: 15.8 ± 1.6; n = 5 control, 5 *Lrp6* embryos; p = 0.5830 by unpaired t-test). Scale bars represent 100 µm; Anterior is up in all images.

**Supplemental Figure 4:**
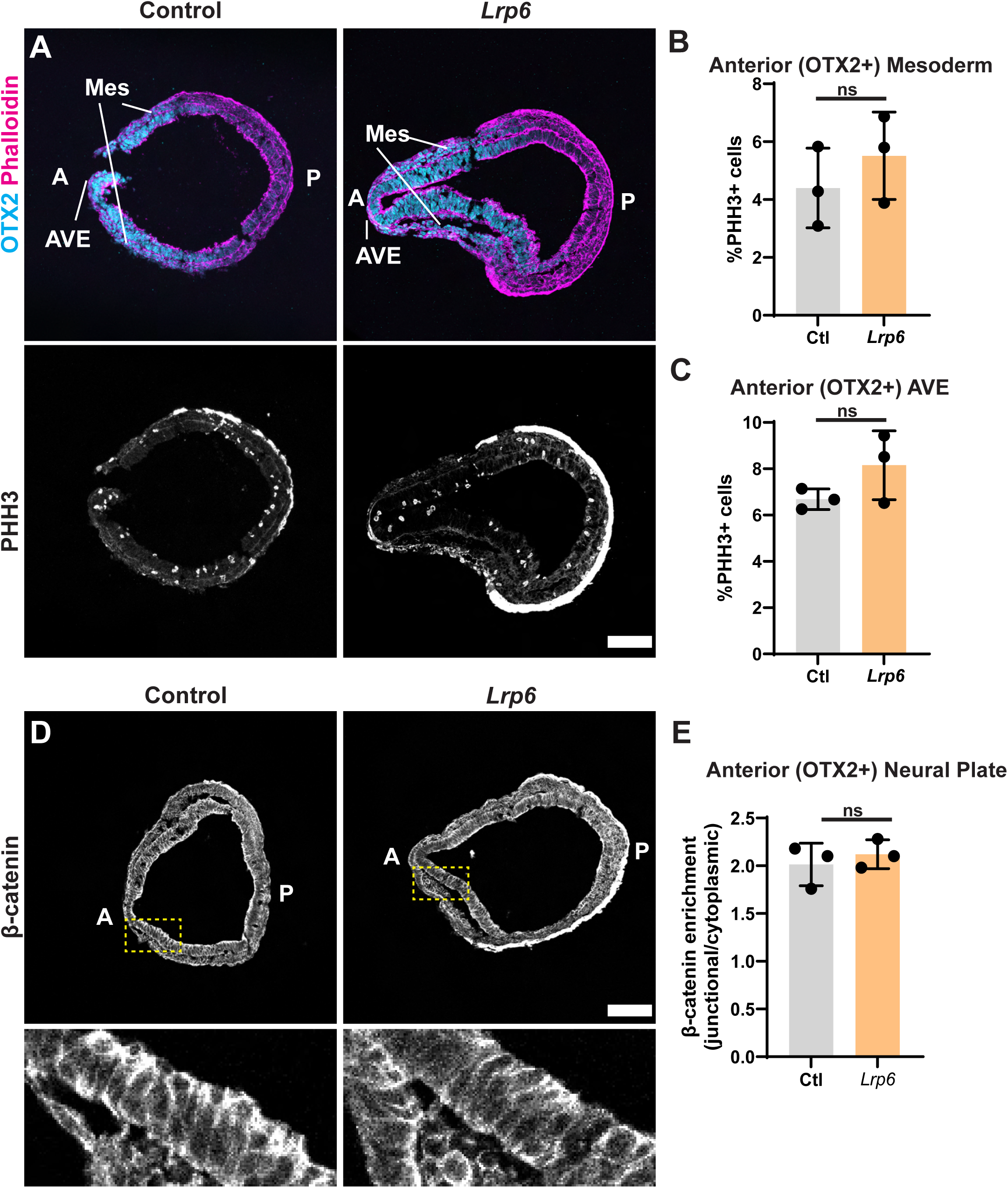
*Lrp6* mutants do not show statistically significant differences in proliferation in non-neural tissues or β-catenin organization at early neural plate stages. (**A**) Transverse sections through control and *Lrp6* mutants at E7.5 stained with an antibody against OTX2 and counterstained with phalloidin (top) or the S-M phase marker phospho-HistoneH3 (bottom). The anterior visceral endoderm (AVE) and mesodermal (Mes) tissues are indicated. (**B**) The ratio of phospho-HistoneH3 positive cells does not significantly differ in the mesoderm (control, 4.40 ± 1.4 % of cells; *Lrp6*, 5.51 ± 1.5 % of cells. n = 3 control and 3 *Lrp6* embryos, p = 0.3984 by unpaired t-test). (**C**) The ratio of phospho-HistoneH3 positive cells does not significantly differ in the anterior visceral endoderm (control, 6.69 ± 0.5 % of cells; *Lrp6*, 8.15 ± 1.5 % of cells. n = 3 control and 3 *Lrp6* embryos, p = 0.1771 by unpaired t-test). (**D**) Transverse sections of E7.5 control and *Lrp6* mutants stained with an antibody against β-catenin (top) and magnified views of the regions highlighted in the dashed yellow boxes (bottom). (**E**) Quantification of the enrichment of junctional β-catenin relative to the cytoplasmic population reveals no difference between control and *Lrp6* mutants (control, 2.01 ± 0.22; *Lrp6*, 2.12 ± 0.15. n = 3 control and 3 *Lrp6* embryos, p = 0.5304 by unpaired t-test). Scale bars represent 100 µm; A, anterior; P, posterior.

**Supplemental Figure 5:**
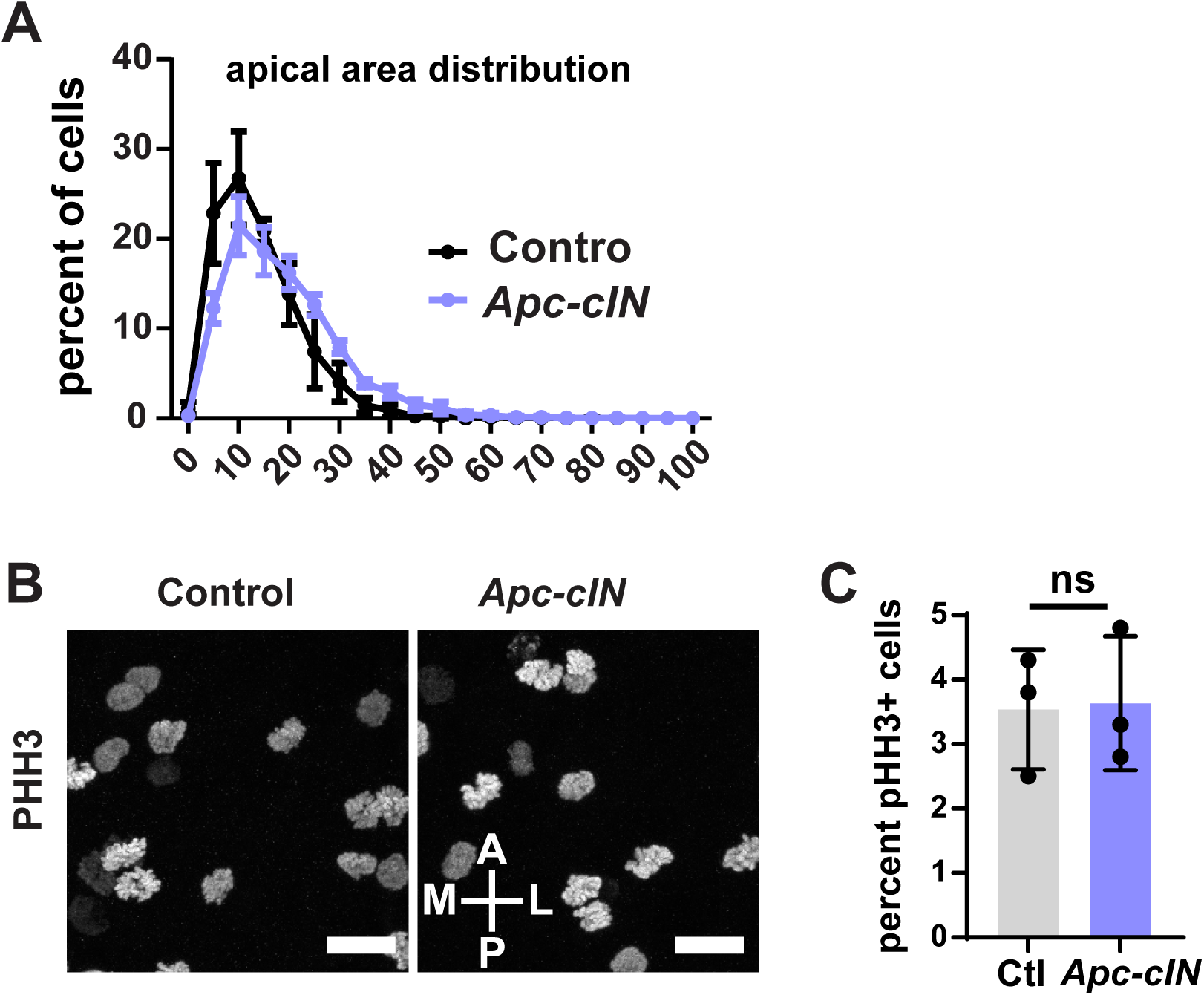
*Apc-cIN* mutants do not show altered proliferative capacity. (**A**) The distribution of apical cell areas in *Apc-cIN* embryos is shifted towards higher areas as compared to controls. (**B**) Images showing staining for phospho-HistoneH 3 in 100 µm x 100 µm lateral regions of control and *Apc-cIN* mutants. (C) Quantification of the proportion of proliferative cells shows no difference between control and *Apc-cIN* mutant embryos (Ctl: 3.53 ± 0.93%; *Apc-cIN*: 3.63 ± 1.04%; n = 3 control, 3 *Apc-cIN*; p = 0.9072 by unpaired t-test). Scale bars represent 20 µm; A, anterior; P, posterior; M, medial; L, lateral.

**Supplemental Figure 6:**
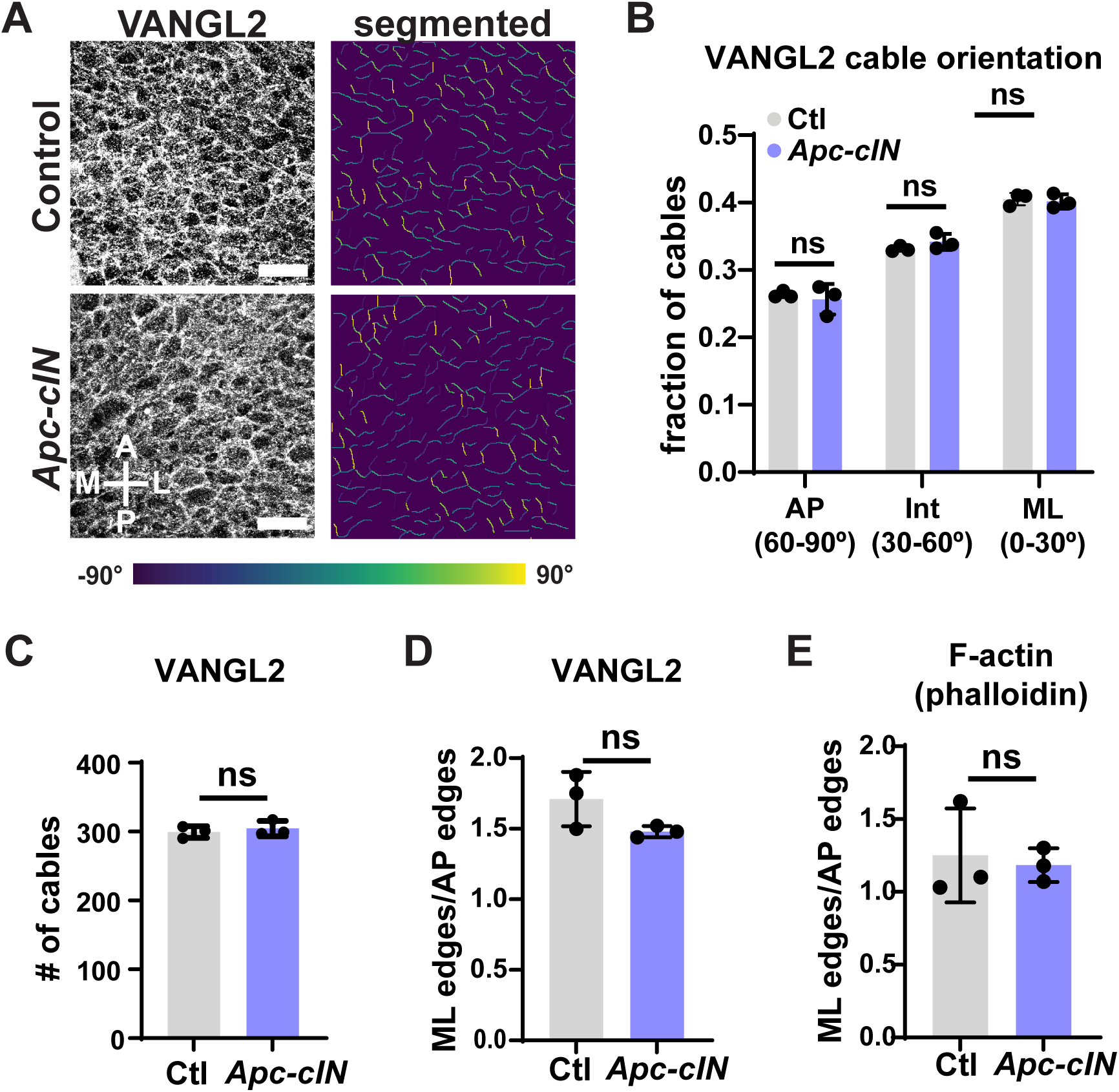
*Apc-cIN* mutants do not show altered planar polarization of VANGL2. (**A**) Staining for the planar cell polarity component VANGL 2 is shown in 100 µm x 100 µm lateral regions (left), and segmented cables that are color coded by their angular orientation (right, 90° is anterior). (**B**) Control and *Apc-cIN* mutants show equivalent planar polarization of VANGL2 staining (two-way ANOVA with Sidak’s multiple comparisons, see Table S1 for n and p values). (**C**) No difference was observed in the number of VANGL2 cables detected in a 100 µm x 100 µm region in control and *Apc-cIN* mutant embryos (Ctl: 299.3 ± 8.6; *Apc-cIN*: 304.3 ± 11.2; n = 3 control, 3 *Apc-cIN*; p = 0.5694 by unpaired t-test). (**D**) The polarized enrichment of VANGL2 mediolaterally did not differ between control and *Apc-cIN* embryos (Ctl: 1.71 ± 0.19; *Apc-cIN*: 1.48 ± 0.04; n = 3 control, 3 *Apc-cIN*; p = 0.1135 by unpaired t-test). (**E**) The polarized enrichment of F-actin mediolaterally did not differ between control and *Apc-cIN* embryos (Ctl: 1.25 ± 0.32; *Apc-cIN*: 1.18 ± 0.12; n = 3 control, 3 *Apc-cIN*; p = 0.7528 by unpaired t-test). Scale bars represent 20 µm; A, anterior; P, posterior; M, medial; L, lateral.

## Table Legends

**Supplemental Table 1: Summary of statistical tests, n, and p values.** For each formal statistical test reported here, the relevant figure, test type, n values, mean and standard deviation (when applicable), and p values are reported.

