## Supplemental Table 1 for "Canonical Wnt pathway modulation is required to correctly execute multiple independent cellular dynamic programs during cranial neural tube closure"

| Figure | Statistical Test | N value | Mean (+/- SD) | P value |
| --- | --- | --- | --- | --- |
| 1B-C | Fisher's Exact Test comparing cranial closure defect penetrance between global (8/10) and Wnt1-Cre2 conditional (0/14) *Lrp6* mutants | 24 | N/A | P < 0.0001 |
| 2C | Two-way ANOVA w/Tukey's multiple comparisons | 5 control embryos per position; 3 *Lrp6* embryos per position | Mean difference<br>20: 0.04349<br>40: 0.05430<br>60: 0.05913<br>80: 0.06906<br>100: 0.08848<br>120: 0.08857<br>140: 0.09426<br>160: 0.08239<br>180: 0.08370<br>200: 0.08370<br>220: 0.1061<br>240: 0.1207<br>260: 0.1503<br>280: 0.1687 | 0: 0.0538<br>20: 0.0154<br>40: 0.0143<br>60: 0.0116<br>80: 0.0021<br>100: 0.0018<br>120: 0.0033<br>140: 0.0035<br>160: 0.0460<br>180: 0.1779<br>200: 0.0767<br>220: 0.1021<br>240: 0.110<br>260: 0.1126<br>280: 0.1557 |

| | | | | |
|---|---|---|---|---|
| 3B, length | Unpaired t-test | 7 control embryos; 6 *Lrp6* embryos | Control: 452.7 +/- 41.87<br>*Lrp6*: 499.1 +/- 76.00 | 0.1905 |
| 3B, width | Unpaired t-test | 7 control embryos; 6 *Lrp6* embryos | Control: 561.4 +/- 40.86<br>*Lrp6*: 713.9 +/- 115.6 | 0.0073 |
| 3D, length | Unpaired t-test | 5 control embryos; 7 *Lrp6* embryos | Control: 1035 +/- 175.7<br>*Lrp6*: 978 98.83 | 0.4845 |
| 3D, width | Unpaired t-test | 5 control embryos; 7 *Lrp6* embryos | Control: 688.6 +/- 108.3<br>*Lrp6*: 1097 100.9 | < 0.0001 |
| 3E, length | Unpaired t-test | 8 control embryos; 6 *Apc-cIN* embryos | Control: 1259 +/- 208.6<br>*Apc-cIN*: 1191 +/- 214.7 | 0.5604 |
| 3F, width | Unpaired t-test | 8 control embryos; 6 *Apc-cIN* embryos | Control: 602.1 +/- 41.09<br>*Apc-cIN*: 616.2 +/- 56.29 | 0.5967 |
| 4D | Unpaired t-test | 4 control embryos; 7 *Lrp6* embryos | Control: 23.63 +/- 4.587<br>*Lrp6*: 24.35 +/- 2.810 | 0.7492 |
| 4F | Unpaired t-test | 6 control embryos; 6 *Lrp6* embryos | Control: 38.96 +/- 6.407<br>*Lrp6*: 40.44 +/- 3.224 | 0.6242 |
| 4H | Unpaired t-test | 4 control embryos; 7 *Lrp6* embryos | Control: 4.115 +/- 0.5507<br>*Lrp6*: 4.109 +/- 0.5454 | 0.9855 |
| 4I | Two-way ANOVA with Sidak's multiple comparisons | 5 control embryos; 5 *Lrp6* embryos | Control AP: 0.5031<br>Control Int: 0.3051<br>Control ML: 0.1918<br>*Lrp6* AP: 0.4776<br>*Lrp6* Int: 0.2978<br>*Lrp6* ML: | AP: 0.9746<br>Int: 0.9994<br>ML: 0.9485 |

| | | | | |
|---|---|---|---|---|
| | | | 0.2245 | |
| 5B | Two-way ANOVA with multiple Sidak's comparisons | 3 control embryos; 3 *Lrp6* embryos | Control AP: 0.2693<br>Control Int: 0.3161<br>Control ML: 0.4089<br>*Lrp6* AP: 0.2808<br>*Lrp6* Int: 0.3225<br>*Lrp6* ML: 0.3979 | AP: 0.9155<br>Int: 0.9812<br>ML: 0.9027 |
| 5C | Unpaired t-test | 3 control embryos; 3 *Lrp6* embryos | Control: 1.74 +/- 0.19<br>*Lrp6*: 1.75 +/- 0.40 | 0.9709 |
| 5E | Two-way ANOVA with Sidak's multiple comparisons | 5 control embryos; 5 *Lrp6* embryos | Control AP: 0.2365<br>Control Int: 0.3340<br>Control ML: 0.4295<br>*Lrp6* AP: 0.2595<br>*Lrp6* Int: 0.3386<br>*Lrp6* ML: 0.4019 | AP: 0.5074<br>Int: 0.9919<br>ML: 0.3570 |
| 5F | Unpaired t-test | 3 control embryos; 3 *Lrp6* embryos | Control: 1.39 +/- 0.18<br>*Lrp6*: 1.43 +/- 0.20 | 0.7915 |
| 6C | Unpaired t-test | 3 control embryos; 3 *Lrp6* embryos | Control: 61.67 +/- 6.028<br>*Lrp6*: 63.67 +/- 3.055 | 0.6352 |
| 6D | Unpaired t-test | 3 control embryos; 3 *Lrp6* embryos | Control: 5.227 +/- 0.9703<br>*Lrp6*: 9.380 +/- 1.251 | 0.0105 |
| 6E | Unpaired t-test | 3 control embryos; 3 *Lrp6* embryos | Control: 5.743 +/- 2.082<br>*Lrp6*: 6.333 +/- 0.9504 | 0.6783 |

| | | | | |
|---|---|---|---|---|
| 7C (length) | Unpaired t-test | 8 control embryos; 6 *Apc-cIN* embryos | Control: 1259 +/- 208.6<br>*Lrp6*: 1191 +/- 214.7 | 0.5604 |
| 7C (width) | Unpaired t-test | 8 control embryos; 6 *Apc-cIN* embryos | Control: 602.1 +/- 41.1<br>*Lrp6*: 616.2 +/- 56.3 | 0.5967 |
| 8B | Unpaired t-test | 4 control embryos; 4 *Apc-cIN* embryos | Control: 1.77 +/- 0.43<br>*Lrp6*: 2.10 +/- 0.26 | 0.2348 |
| 8D | Unpaired t-test | 4 control embryos; 4 *Apc-cIN* embryos | Control: 2.55 +/- 0.27<br>*Lrp6*: 1.88 +/- 0.08 | 0.0030 |
| 9G | Unpaired t-test | 3 control embryos; 3 *Apc-cIN* embryos | Control: 13.96 +/- 2.286<br>*Apc-cIN*: 18.78 +/- 1.074 | 0.0293 |
| 9H | Unpaired t-test | 3 control embryos; 3 *Apc-cIN* embryos | Control: 50.61 +/- 5.772<br>*Apc-cIN*: 51.10 +/- 5.924 | 0.9238 |
| 9I (apical) | Unpaired t-test | 4 control embryos; 4 *Apc-cIN* embryos | Control: 3.57 +/- 0.35<br>*Apc-cIN*: 5.52 +/- 0.99 | 0.0100 |
| 9I (basal) | Unpaired t-test | 4 control embryos; 4 *Apc-cIN* embryos | Control: 2.37 +/- 0.28<br>*Apc-cIN*: 2.46 +/- 0.51 | 0.7765 |
| 9J (junctions) | Unpaired t-test | 4 control embryos; 4 *Apc-cIN* embryos | Control: 80.44 +/- 27.96<br>*Apc-cIN*: 73.01 +/- 11.22 | 0.6392 |
| 9J (medio-apical) | Unpaired t-test | 4 control embryos; 4 *Apc-cIN* embryos | Control: 28.10 +/- 8.10<br>*Apc-cIN*: 16.80 +/- 3.42 | 0.0422 |
| S1B | Unpaired t-test | 4 control embryos; 4 *Lrp6* embryos | Control: 1.79 +/- 0.23<br>*Lrp6*: 1.82 +/- 0.10 | 0.8048 |
| S1C | Unpaired t-test | 4 control embryos; 4 *Lrp6* embryos | Control: 4.79 +/- 0.72<br>*Lrp6*: 4.20 +/- 0.47 | 0.2185 |

| | | | | |
|---|---|---|---|---|
| S1E | Unpaired t-test | 4 control embryos; 4 *Lrp6* embryos | Control: 2.76 +/- 0.32<br>*Lrp6*: 2.72 +/- 0.08 | 0.9225 |
| S1F | Unpaired t-test | 4 control embryos; 4 *Lrp6* embryos | Control: 3.06 +/- 0.63<br>*Lrp6*: 2.55 +/- 0.44 | 0.2369 |
| S2A | Two-way ANOVA with Sidak's multiple comparisons | 5 control embryos; 3 *Lrp6* embryos | Control AP: 0.2622<br>Control Int: 0.3308<br>Control ML: 0.4070<br>*Lrp6* AP: 0.2743<br>*Lrp6* Int: 0.3409<br>*Lrp6* ML: 0.3848 | AP: 0.9566<br>Int: 0.9742<br>ML: 0.7915 |
| S2B | Two-way ANOVA with Sidak's multiple comparisons | 5 control embryos; 3 *Lrp6* embryos | Control AP: 0.2591<br>Control Int: 0.3046<br>Control ML: 0.4363<br>*Lrp6* AP: 0.3161<br>*Lrp6* Int: 0.2661<br>*Lrp6* ML: 0.4178 | AP: 0.3041<br>Int: 0.6207<br>ML: 0.9343 |
| S2C | Two-way ANOVA with Sidak's multiple comparisons | 5 control embryos; 3 *Lrp6* embryos | Control AP: 0.2960<br>Control Int: 0.3245<br>Control ML: 0.3795<br>*Lrp6* AP: 0.2305<br>*Lrp6* Int: 0.3810<br>*Lrp6* ML: 0.3885 | AP: 0.1197<br>Int: 0.2071<br>ML: 0.9874 |
| S2D | Two-way ANOVA with Sidak's | 5 control embryos; 3 *Lrp6* embryos | Control AP: 0.2347 | AP: 0.9691<br>Int: 0.3193<br>ML: 0.5580 |

| | | | | |
|---|---|---|---|---|
| | multiple comparisons | | Control Int: 0.3541<br>Control ML: 0.4112<br>*Lrp6* AP: 0.2504<br>*Lrp6* Int: 0.2916<br>*Lrp6* ML: 0.4580 | |
| S2E | Unpaired t-test | 5 control embryos; 3 *Lrp6* embryos | Control: 229 +/- 28.59<br>*Lrp6*: 234.3 +/- 45.98 | 0.8432 |
| S2F | Unpaired t-test | 5 control embryos; 3 *Lrp6* embryos | Control: 258.6 +/- 33.72<br>*Lrp6*: 309.0 +/- 63.32 | 0.1823 |
| S3C | Unpaired t-test | 3 control embryos; 3 *Lrp6* embryos | Control: 120.5 +/- 8.81<br>*Lrp6*: 194.8 +/- 16.36 | <0.0001 |
| S3D | Unpaired t-test | 5 control embryos; 5 *Lrp6* embryos | Control: 16.40 +/- 1.67<br>*Lrp6*: 15.80 +/- 1.64 | 0.5830 |
| S4B | Unpaired t-test | 3 control embryos; 3 *Lrp6* embryos | Control: 4.400 +/- 1.374<br>*Lrp6*: 5.513 +/- 1.511 | 0.3984 |
| S4C | Unpaired t-test | 3 control embryos; 3 *Lrp6* embryos | Control: 6.687 +/- 0.445<br>*Lrp6*: 8.153 +/- 1.487 | 0.1771 |
| S4E | Unpaired t-test | 3 control embryos; 3 *Lrp6* embryos | Control: 2.01 +/- 0.22<br>*Lrp6*: 2.12 +/- 0.15 | 0.5304 |
| S5A | Kolmogorov-Smirnov test*<br><br>*Requires collapsing replicates into a single ctl distribution and a single *Apc-cIN* distribution, may | 3309 Control cells, 2449 *Apc-cIN* cells | Ctl: 13.75 +/- 8.448<br>*Apc-cIN*: 18.76 +/- 11.03 | <0.0001 |

| | | | | |
|---|---|---|---|---|
| | inappropriately overpower test | | | |
| S5C | Unpaired t-test | 3 control embryos; 3 *Apc-cIN* embryos | Control: 3.533 +/- 0.9292<br>*Apc-cIN*: 3.633 +/- 1.041 | 0.9072 |
| S6B | Two-way ANOVA with Sidak's multiple comparisons | 3 control embryos; 3 *Apc-cIN* embryos | Control AP: 0.2635<br>Control Int: 0.3312<br>Control ML: 0.4052<br>*Apc-cIN* AP: 0.2566<br>*Apc-cIN* Int: 0.3417<br>*Apc-cIN* ML: 0.4017 | AP: 0.8737<br>Int: 0.6782<br>ML: 0.9807 |
| S6C | Unpaired t-test | 3 control embryos; 3 *Apc-cIN* embryos | Control: 299.3 +/- 8.622<br>*Apc-cIN*: 304.3 +/- 11.02 | 0.5694 |
| S6D | Unpaired t-test | 3 control embryos; 3 *Apc-cIN* embryos | Control: 1.71 +/- 0.19<br>*Apc-cIN*: 1.48 +/- 0.04 | 0.1135 |
| S6E | Unpaired t-test | 3 control embryos; 3 *Apc-cIN* embryos | Control: 1.25 +/- 0.32<br>*Apc-cIN*: 1.18 +/- 0.12 | 0.7528 |
